# Tensor Image Registration Library: Automated Non-Linear Registration of Sparsely Sampled Histological Specimens to Post-Mortem MRI of the Whole Human Brain

**DOI:** 10.1101/849570

**Authors:** Istvan N. Huszar, Menuka Pallebage-Gamarallage, Sean Foxley, Benjamin C. Tendler, Anna Leonte, Marlies Hiemstra, Jeroen Mollink, Adele Smart, Sarah Bangerter-Christensen, Hannah Brooks, Martin R. Turner, Olaf Ansorge, Karla L. Miller, Mark Jenkinson

**Affiliations:** Wellcome Centre for Integrative Neuroimaging, FMRIB, Nuffield Department of Clinical Neurosciences, University of Oxford, Oxford, UK; Nuffield Department of Clinical Neurosciences, University of Oxford, Oxford, UK; Department of Radiology, University of Chicago, Chicago, IL, USA; Department of Neuroscience, University of Groningen, Groningen, Netherlands; Department of Anatomy, Donders Institute for Brain, Cognition and Behaviour, Radboud University Medical Centre, Nijmegen, Netherlands; Brigham Young University, Provo, UT, USA

**Keywords:** registration, histology, post-mortem, MRI, brain, human

## Abstract

There is a need to understand the histopathological basis of MRI signal characteristics in complex biological matter. Microstructural imaging holds promise for sensitive and specific indicators of the early stages of human neurodegeneration but requires validation against traditional histological markers before it can be reliably applied in the clinical setting. Validation relies on a precise and preferably automatic method to align MRI and histological images of the same tissue, which poses unique challenges compared to more conventional MRI-to-MRI registration.

A customisable open-source platform, Tensor Image Registration Library (TIRL) is presented. Based on TIRL, a fully automated pipeline was implemented to align small stained histological images with dissection photographs of corresponding tissue blocks and coronal brain slices, and further with high-resolution (0.5 mm) whole-brain post-mortem MRI data. The pipeline performed three separate deformable registrations to achieve accurate mapping between whole-brain MRI and small-slide histology coordinates. The robustness and accuracy of the individual registration steps were evaluated using both simulated data and real-life images from 6 different anatomical locations of one post-mortem human brain.

The automated registration method demonstrated sub-millimetre accuracy in all steps, robustness against tissue damage, and good reproducibility between experiments. The method also outperformed manual landmark-based slice-to-volume registration, also correcting for curvatures in the slicing plane. Due to the customisability of TIRL, the pipeline can be conveniently adapted for other research needs and is therefore suitable for the large-scale comparison of routinely collected histology and MRI data.

**Highlights:**

- TIRL: new framework for prototyping bespoke image registration pipelines
- Pipeline for automated registration of small-slide histology to whole-brain MRI
- Slice-to-volume registration accounting for through-plane deformations
- No need for serial histological sampling

## 1. Introduction

### 1.1. Motivation

Histopathological studies have contributed an essential part to our understanding of human neurodegeneration. Looking at a chemically stained post-mortem tissue sample under a microscope, one can find molecular evidence for whether or not the observed region of the central nervous system has been affected by a disease process. Protein aggregates and neuronal death are among the defining histological features of neurodegeneration [1], usually predating clinical symptoms by several years [2]. Post-mortem studies of Parkinson’s disease [3], Alzheimer’s disease [4] and amyotrophic lateral sclerosis (ALS) [5] have indicated that the type of aggregates and their spatial distribution in the central nervous system are together representative of the type of neurodegeneration. Hence the concept of neurodegeneration as a prion-like spatiotemporal process has emerged [6, 7].

Being restricted to post-mortem tissue, histology alone provides limited information about the temporal aspect of the disease and it is often used to study certain regions only instead of probing the whole human brain. Intra-individual characterisation of neurodegeneration as a spatiotemporal process therefore requires the combination of histology with a suitable in-vivo imaging technique that provides full brain coverage, is repeatable and desirably harmless for the patient.

Advanced magnetic resonance imaging (MRI) techniques, in addition to capturing gross 3D anatomical and functional images of the whole brain, can also interrogate tissue properties at microscopic scales far smaller than the resolved voxel size. When applied to the human brain, these emerging microstructural MRI methods aim to estimate tissue properties such as neurite density, intracellular volume fraction [8], axon diameter [9], myelin content [10, 11], and cortical fibre orientation [12] usually by interpreting local changes of the MRI signal in the framework of sophisticated biophysical models [13] of tissue structure. Collectively these type of methods have been regarded as “in-vivo histology” [14, 15] and serve as a promising non-invasive tool for tracking tissue-level spatiotemporal changes related to human neurodegeneration. However, it is a matter of active debate [16] how the measured quantities relate to actual histological parameters, especially in disease.

In order to characterise the relationship between MRI signal alterations and histopathological features in motor neuron disease (MND), our group previously acquired multi-modal MRI scans of whole, post-mortem human brains that were subsequently dissected into blocks for histopathological staining [17]. This dataset is the subject of an ongoing research project and will be released in full upon its completion. In the present paper, we address how the resultant 2D histological images can be accurately registered to the 3D whole-brain post-mortem MRI data in an automated way, enabling systematic voxel-wise comparison between MRI and histological parameters. First, we provide an overview of existing approaches to MRI–histology registration, then describe the development of a novel open-source image registration framework that we used to successfully register images from our dataset.

### 1.2. Previous work in MRI-histology registration

It is important to distinguish between two main approaches to MRI-histology registration based on how the histology data is collected, as it largely determines how the registration is performed. Over the next few paragraphs we shortly review previously proposed registration methods for (1) dense systematic histological sampling, and (2) stand-alone histological images.

Methods of the first kind are well-developed with numerous examples [18-28] from as early as 1994. A comprehensive review of the methods in this category can be found in *Pichat* et al [29]. For these methods, tissues must be frozen or embedded in a rigid medium and sectioned at regular intervals. Most commonly the tissue block face is photographed after each section to serve as a rigid reference. Distortions of the thin tissue sections are compensated by 2D deformable registration to the corresponding rigid reference, which are subsequently stacked to create a volume of photographic/histological data. This volume is later registered to the MRI data using standard 3D registration tools such as ABA [30] or ANTs [31]. As a novelty, *Iglesias* et al [32] recently demonstrated accurate (0.5 – 2 mm) 2D-to-2D histology-to-MRI registration without photographic intermediates. Assuming perfect slice correspondence, they mapped sequentially sampled whole-brain histology images to an MRI volume. Complementary to this work, *Pichat* et al [33] proposed a method for direct histology-to-MRI registration between small histological samples (as opposed to whole-hemisphere images) and corresponding MRI slices via automated affine-invariant shape matching, but only reported preliminary results that required manual MRI slice matching.

Perhaps the biggest advantage of the methods in this category is that the 3D geometrical correspondence of the sections is accurately preserved. With the use of a rigid reference, the “banana effect”, [34] in which an accumulation of small shifts between adjacent slices results in a shearing in the third dimension, may also be avoided. However, given the size of the human brain, the acquisition of systematically sampled histology data requires bespoke slicing and stain automation hardware, none of which is readily available at most neuropathology facilities. This approach is therefore better suited to study the brains of small animals or, with substantial time and workforce commitment, a single human brain [35, 36].

For stand-alone histological images (i.e. a single, small-format slide), a direct slice-to-volume registration must be employed. Despite the fact that almost the entire body of histology images that have ever been created in neuropathology facilities for expert interpretation exist in this format, suitable registration methods are disproportionately underrepresented in the literature. This might be due to the unique challenge associated with slice-to-volume registration: the parameters that are necessary to align a potentially distorted 2D image in 3D space have a vast search space, and a high propensity to converge to local optima, as a small 2D slice may constitute a relatively good fit at many locations in 3D space. Most reported pipelines are semi-automatic, requiring accurate manual slice initialisation or annotating correspondent anatomical structures.

*Kim* et al [37] reported the first relevant slice-to-volume registration approach between *post-mortem* brain slice photographs and *ante-mortem* MRI, using 2^nd-^ and 3^rd^-order polynomial parametrisation for in-plane and out-of-plane deformations. The method was later used by *Singh* et al [38] to register histological images of vascular lesions to in-vivo MRI data using photographic intermediates, reporting an overall registration accuracy of 3-8 mm. *Meyer* et al [39] reported a semi-automated registration method to align a histological section of murine glioma to in-vivo MRI using high-resolution *ex-vivo* MRI as an intermediate reference volume. As an improvement to using polynomials to parametrise deformations, they used thin-plate splines (TPS) to correct for both in-plane and out-of-plane deformations of the histological section. However, the accuracy of their method was not mentioned, and the code was not published. *Osechinskiy* et al [40] registered histological sections of whole hemispheres to 3D MRI data and conducted a comprehensive analysis of cost functions, optimisation and transformation methods. The best results were achieved by using TPS-based parametrisation of a 3D deformation field, optimised by the NEWUOA algorithm [41] for Pearson’s correlation of the images. As a further refinement to the technique, they devised a novel similarity metric based on the correspondence of grey-white matter boundaries [42]. Unfortunately, the authors never made their implementation publicly available for reuse, prohibiting the test of novel texture-based cost functions, such as the Modality-Independent Neighbourhood Descriptor (MIND) [43]. The registration of small histological sections (as opposed to whole-hemisphere images) was studied by *Ohnishi* et al [44]. Using manual landmarks, they stitched together smaller histological images into a full hemisphere, and subsequently registered it to a photograph of the coronal section of the hemisphere, which was further registered to 3D MRI. Neither in-plane nor out-of-plane deformations were considered in the slice-to-volume step; the authors instead recommended using specialised hardware to avoid distortions.

In both categories of problems described above, the pipelines mentioned so far were built for a specific purpose and have not been released in the form of open-source software. As a consequence, new experimenters are repeatedly required to invest time and effort into extensive hardware and/or software development [45], which negatively impacts large-scale validation studies. Recently, *Majka* et al [46] released Possum, an open-source framework for reconstructing volumetric histology data, which is a great step toward standardising this aspect of stack-based MRI-histology registration. A recent preprint by *Alegro* et al [36, 47] reported an automated pipeline for serial histological sections. To the best of our knowledge, a similar software tool for slice-to-volume registration of small stand-alone histological images to MRI data is still not available to date.

In the present work, we aim to address this need by describing a new open-source image registration framework that aims to integrate previously published methods in a single package, providing a customisable workflow that is compatible with most common image formats for MRI, histology and photographs. Based on this framework, we propose a fully automated registration pipeline to align small histological sections with volumetric MRI data using photographic intermediates. As an additional novelty, we demonstrate by example that in the case of free-hand brain cutting, involuntary deflections from the slicing plane are large enough to disrupt the anatomical correspondence between histological sections and visually matched slices of the MRI volume, and explain how these distortions are compensated within the proposed pipeline.

The organisation of the paper is as follows. In ‘Materials and Methods’ we describe the acquisition (*section 2.1*) and pre-processing (*section 2.2*) of the imaging data, formulate the registration problem (*section 2.4*), introduce the Tensor Image Registration Library (TIRL) (*section 2.5*), and finally describe each stage of the proposed MRI-histology registration pipeline (*sections 2.6-2.8*) as well as the combination of all stages (*section 2.9*). In ‘Results’ we present summative or representative registration results and describe the accuracy of each stage (*sections 3.1-3.3*) as well as showing an example of end-to-end MRI-histology registration (*section 3.4*). Finally, in *section 3.5* we introduce an optional stage that may be used to refine end-to-end registration results and show a further example of MRI-histology alignment. In the ‘Discussion’ section we highlight potential directions for further improvement and finally identify the role of the current developments in the broader context of neuroimaging research.

## 2. Material and Methods

### 2.1. Imaging data acquisition

Figure 1 summarises the collection of the imaging data that served as a starting point for the present study. All data was collected and used according to the Oxford Brain Bank’s (OBB) generic Research Ethics Committee approval (15/SC/0639). Written informed consent was obtained by the OBB from all participants of this study. Thirteen formalin-fixed post-mortem brains with neuropathologically and clinically verified MND were obtained from the cases of the OBB between 2014 and 2017. The median age of the donors was 65.5 years at death (full range: 27-77 years), and ten of them were males, two of them females. An additional three brains with no neuropathological hallmarks or clinical records of neurodegeneration were obtained as controls (age at death: 61, 76, 89 years; 2 males, 1 female). The post-mortem interval of the brains varied between 1 and 7 days (median: 3 days). The brains (denuded of the dura mater) were immersed in 10% neutral buffered formalin for at least 1 month to allow even preservation of the tissues throughout the full volume of the brain. Before scanning, each specimen was placed into a brain-shape plastic container to prevent large deformations and the container was filled with Fluorinert to supress the background signal. Scans were performed on a clinical 7T Siemens MRI scanner at the Wellcome Centre for Integrative Neuroimaging (University of Oxford) using an optimised 48-hour acquisition protocol yielding quantitative T1 and T2 maps at 1 mm isotropic resolution, T2* and susceptibility maps at 0.5 mm isotropic resolution, DWI at 0.85 mm isotropic resolution and a bSSFP anatomical reference scan at 0.25 mm isotropic resolution (also referred to as 3D-TRUFI) [17, 48]. The MRI images were aligned and post-processed with an in-house pipeline (*B.C. Tendler*, in preparation). For the purposes of the present paper, only the anatomical reference scan (resampled to 0.5 mm isotropic resolution) was used, because it exhibited the highest contrast between grey and white matter.

**Figure 1.**
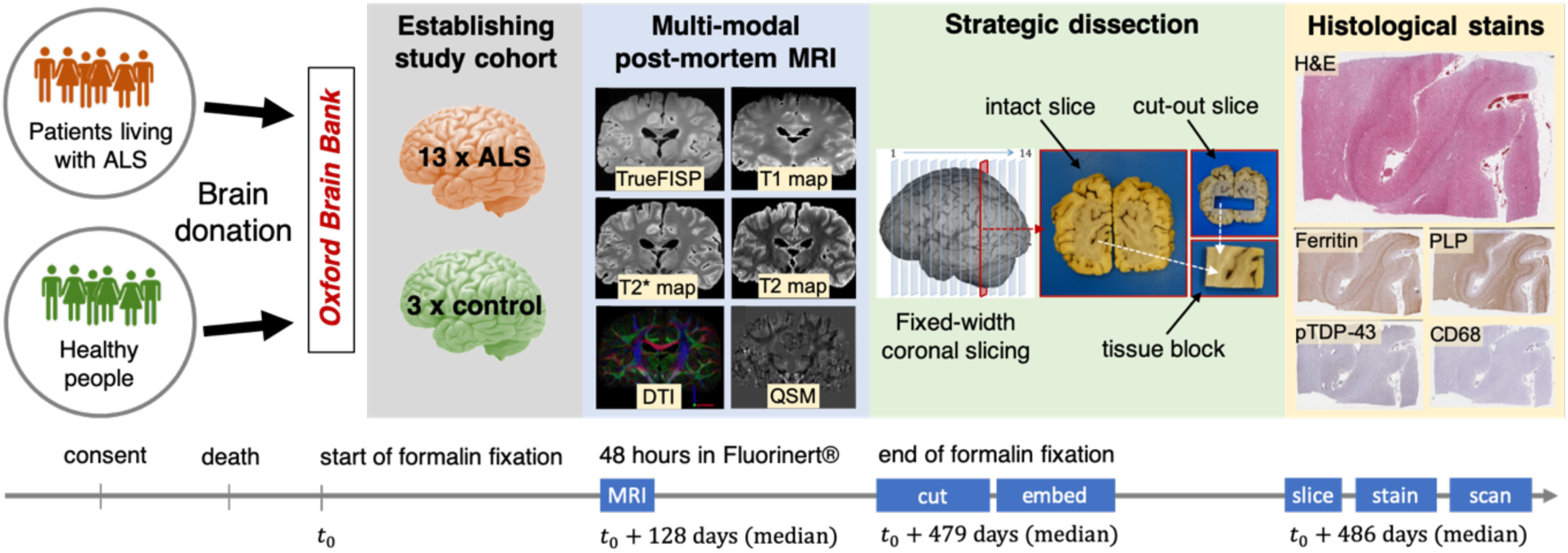
Overview of image data collection. Whole human brains were obtained from the Oxford Brain Bank from 3 consented healthy individuals and 13 patients with MND. Multi-modal quantitative MR images were acquired from each brain after at least 1 month of formalin fixation (4 months on average). The brains were dissected with a standard protocol for histopathological staging. Coronal brain slices were photographed on both sides before and after the excision of smaller tissue blocks of interest, which were also photographed. H&E, immunostains for ferritin, PLP, pTDP-43, and CD68 stains were created from the superficial layers of each tissue block.

The formalin-fixed whole brains were subsequently dissected at the Neuropathology Department of the John Radcliffe Hospital (Oxford). Following an optimised whole-brain sampling protocol [17], the brain was manually sliced into approximately 1 cm thick coronal sections, starting from the plane of the mamillary bodies toward the anterior and posterior poles of the brain. The total number of slices (13-17) varied with the size of the brain. As part of the protocol, *en bloc* dissection of the hand knob from the primary motor (M1) and primary sensory (S1) cortices in advance of the coronal slicing resulted in bilateral damage in a few of the middle slices. (We will refer to the extracted block later as the “M1S1 block”.) From predefined anatomical locations in each slice, one or more smaller tissue blocks were extracted by knife section. The size and shape of the blocks varied across sampling sites, but most of them were not larger than a few centimetres. The whole process was carefully documented by routinely capturing photographs of (1) both the anterior and posterior surfaces of the coronal slices, (2) each coronal slice after the extraction of a new tissue block, and (3) both the anterior and posterior faces of all tissue blocks (Figure 1). The size of the raw photographs was 5472 × 3648, and their resolution was later recorded from a size guide in the images as ∼55 μm/pixel.

Finally, all tissue blocks were embedded in paraffin and sectioned on a microtome at 6-10 μm thickness. Tissue sections were stained by various histological methods, including standard haematoxylin and eosin (H&E) and immunochemistry for proteins of interest such as ferritin, myelin proteolipid protein (PLP), activated microglia and macrophages (CD68), phosphorylated TAR-DNA binding protein-43 (pTDP-43) and pan microglia (Iba-1). Specifically for the purpose of registration with MRI, additional LFB+PAS (Luxol fast blue combined with the periodic acid-Schiff procedure) staining was performed on two blocks of a single brain. Digital whole-slide images were created in SVS format using an Aperio ScanScope slide scanner at ×20 objective magnification, yielding a typical matrix size of approximately 60000 × 45000 at full resolution (∼0.5 μm/pixel).

### 2.2. Image pre-processing

Before entering the registration pipeline, all of the above-mentioned images underwent a number of pre-processing steps to (1) reduce some of the variability of the input and (2) aid registration by addressing structural discrepancies between corresponding images.

#### Ad 1

To standardise the input of the registration pipeline, brain slices and tissue blocks were isolated from other objects in the photographs by cropping the central 50% and 30% of the original images along each axis, and segmented from the blue matte background using *k*-means classification (*k* = 2) of the RGB vectors in the cropped image. The images were smoothed with the mean shift algorithm (*r*_*spatial*_: 3 px, *r*_*RGB*_: 10) before the classification to prevent noisy segmentation within the tissue. Tissue debris and glare occasionally resulted in false positive segmentation that were successfully removed by searching for connected components in the segmented image and discarding anything under an area of 2000 pixels (resolution: 50 μm/px). As a result of the pre-processing, brain slice and tissue block photographs had an approximate size of 2500 × 2500 pixels and 800 × 800 pixels, respectively, 3 colour channels and zero-filled background. The histological images were imported from the lowest resolution level (∼5 μm/px) of the digitised whole-slide images, and further subsampled to match the resolution of the photographs. As suggested by *Jenkinson* and *Smith* [49, 50] Gaussian smoothing was applied to the images before downsampling (with FWHM in mm set to the downsampling factor) to ensure that new pixel values are representative of all pixels in the original image. Finally, all photographs and the histological images were flattened into 8-bit grayscale images by taking the Euclidean norm of the RGB vectors.

#### Ad 2

Due to the post-mortem nature of the study, full anatomical correspondence may not be guaranteed between corresponding image pairs. For example, as long as the cerebellum is removed at the start of the dissection process, coronal sections of the MR volumes will be different from the corresponding autopsy photographs, and lead to severe registration error in occipital slices. Similarly, missing parts of the motor and sensory cortices has a similar consequence for the slices that are close to the centre. These structural discrepancies were also addressed by pre-processing. We used the cerebral segmentation tool in BrainSuite (ver. 18) [51] to perform brain extraction and remove the cerebellum from the high-resolution structural MRI scan before it was fed into the pipeline. Furthermore, hand-drawn binary 2D masks were used to facilitate slice-to-volume registration at the centre of the brain, where parts of the hemispheres were absent from the photographs as a result of removing the M1S1 blocks.

### 2.3. Overview of the registration pipeline

As shown in Figure 2, the proposed automated registration pipeline imports the pre-processed images (histology, photograph, MRI), and performs three consecutive registrations (stages 1, 2, and 3) to map the pixels of a histological image (*x* ∈ ℝ^2^) on the voxels of MRI data (*x*′ ∈ ℝ^3^). An optional extension to the pipeline (stage 4) will be described in *section 3.5*, following the discussion of registration results based on the three main stages.

**Figure 2.**
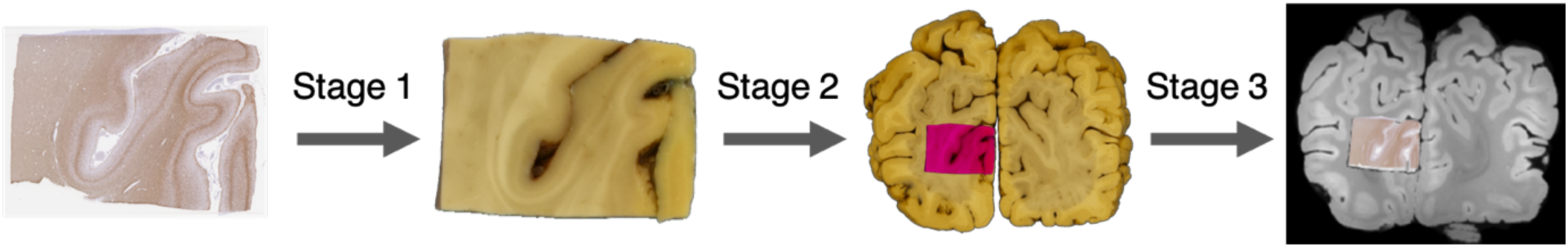
Overview of the three independent deformable registration steps of the pipeline. Stage 1: histology to tissue block photograph, stage 2: tissue block to brain slice photograph, stage 3: brain slice photograph to MRI volume. MRI-histology registration is realised by optimising each stage separately and eventually combining all three stages into a single, final transformation. As discussed later in *section 3.5*, an optional 4^th^ stage may be employed to fine tune the alignment of the registered histological section within the MRI volume.

### 2.4. Formulation of image registration

In the following paragraphs we describe the mathematical formulation for registering two-dimensional scalar-valued (single-channel) images. We do this to keep notations as simple as possible, but the derivation can be readily extended for images with three spatial dimensions and/or multiple channels (i.e. vector-, matrix- or tensor-valued pixels or voxels), and the actual implementation follows the general case.

We define a single-channel target (*T*(***x***) ∈ ℝ) and a single-channel source (*H*(***x***′) ∈ ℝ) image as continuous functions on finite Euclidean domains Ω ⊂ ℝ^*d*^, and Ψ ⊂ ℝ^*d*^ of dimension *d* = 2. We further define a bijection *ϕ*_***p***_(·) with parameters ***p*** that maps the coordinates of corresponding pixels between the source and the target domain: *ϕ*_***p***_: Ψ → Ω, ***x*** = *ϕ*_***p***_(***x***′), and its inverse such that 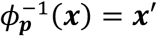. Using this notation, the registration problem between two images may be formalised as:

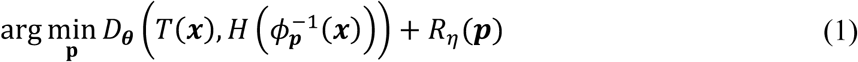

where *D*_*θ*_(·) is a distance function (or cost) with parameters *θ*, that quantifies the dissimilarity of corresponding pixels. *R*_*η*_ is the regularisation term that imposes constraints on the transformation parameters (such as spatial smoothness or elasticity *etc* [52]) and smooths the objective function for efficient optimisation. The most common choice for *D*_*θ*_(·) is the sum of squared intensity differences (SSD):

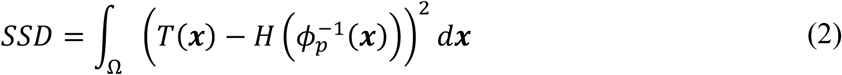

However, SSD becomes problematic when images of different modalities are concerned (such as MRI, CT, photographs, histology *etc.*), as in this case the images may exhibit different internal contrast and the assumption of a monotonic relationship between intensity difference and anatomical dissimilarity no longer prevails. Alternative distance measures that can be used in the multi-modal context are correlation ratio (CR), and normalised mutual information (NMI) [49]. In the context of MRI-histology registration none of these statistical measures are ideal, because the region of interest that corresponds to a histological section may only constitute a small number of voxels in MRI space. In this work, we therefore use a more recently proposed pattern-based approach called the Modality-Independent Neighbourhood Descriptor (MIND) [43], which is a non-linear image operation that enables us to subsequently use SSD on multi-modal data. In essence, MIND captures the local self-similarity of the image by replacing each pixel value with a vector, the components of which describe the intensity relationship of the current pixel with that of its neighbours in a directionally dependent manner.

To calculate the MIND-representation of a single-channel image, we first discretise our previous image definitions such that ***x*** denotes target pixel coordinates (non-negative integers), and we define a small set of neighbourhood intensities around each pixel (Figure 3A, red tiles):

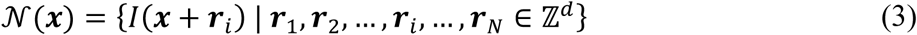

**Figure 3.**
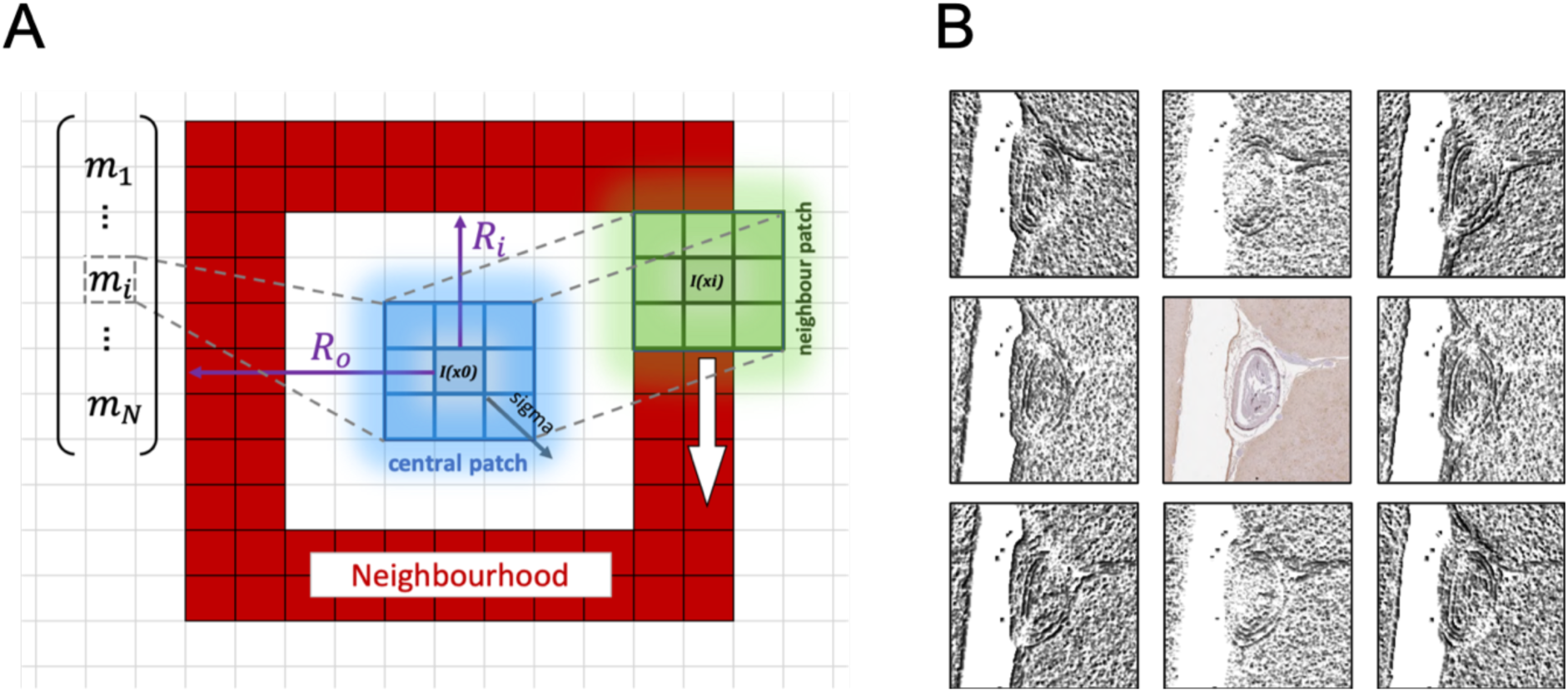
The modality-independent neighbourhood descriptor (MIND). **(A)** schematic of MIND calculation for the pixel at the centre (at ***x***_0_). A neighbourhood (red) is defined as a hollow square plate by the half side lengths of its inner (*R*_*i*_) and outer (*R*_*o*_) bounding squares. Each component of the MIND vector (*m*_*i*_) represents the dissimilarity between the central (blue) patch and the peripheral (green) patch that is centred on the *i*-th element of the neighbourhood (at ***x***_*i*_). Dissimilarity is calculated as the sum of squared differences between corresponding elements of the two patches. Resultant values are normalised by the local variance and taken as a negative exponent of an exponential function (cf. equations in *section 2.4*). Values of the raw MIND vector are finally scaled such that the largest MIND component at each pixel becomes 1. **(B)** Result of the 2D MIND operation (*R*_*i*_ = 0, *R*_*o*_ = 1) on a small portion of an actual histological image. (The image was flattened to grayscale before the MIND operation.) Black-and-white images on the periphery correspond to individual components of the MIND vectors according to their position in the 8-neighbourhood. Note the direction-sensitive enhancement of image edges in each tile.

The neighbourhood may be of arbitrary shape. Here we parametrise it as a hollow square plate that is defined by the half side lengths of its inner (*R*_*i*_ ≥ 0) and outer (*R*_*o*_ ≥ 1, and *R*_*o*_ > *R*_*i*_) bounding squares (see Figure 3A).

In a way that is similar to the definition of the pixel’s neighbourhood, we further define identical patches around both the central pixel (*I*(***x***) ∈ ℝ) (*𝒫*_*c*_) and each of the pixels in its neighbourhood (*𝒫*_1..*i*..*N*_) (Figure 3A, *blue and green squares*):

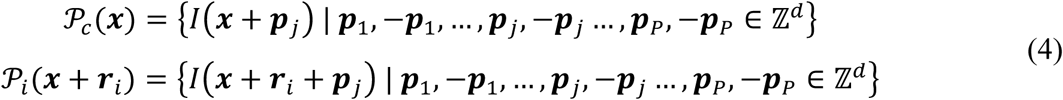

With each of the |*𝒩*| components of the MIND vector the aim is to represent the similarity of the central patch to one of the neighbourhood patches, hence the *i*-th component of the MIND vector ***m***(***x***) at pixel ***x*** is defined as:

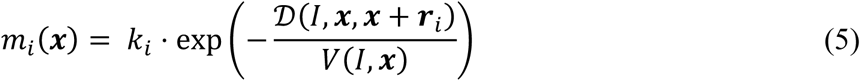

where *𝒟* is the patch-based image dissimilarity index of the image at ***x*** ∈ ℝ^*d*^ with respect to the *i*-th neighbourhood point at ***x*** + ***r***_*i*_. The dissimilarity index is essentially the squared difference between the intensities at ***x*** and ***x*** + ***r***_*i*_, but instead it is calculated as a sum of all squared differences between the corresponding elements of the central and the neighbourhood patches (*𝒫*) to maintain robustness against image noise (Figure 3A, blue and green squares):

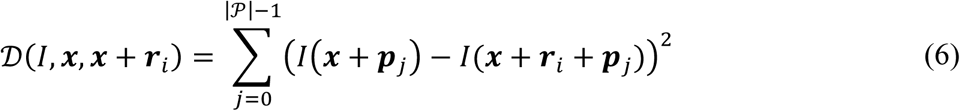

and *V* is the local intensity variance of the image at pixel ***x***:

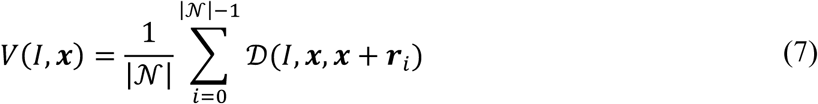

*k*_*i*_ is a normalisation factor that ensures that the largest component of the MIND vector (of size |*𝒩*|) at every ***x*** is 1. As a result of the MIND transformation, the similarity of multi-modal images can be defined as the squared Euclidean distance between the corresponding MIND vectors of the target (***m***_*T*_(***x***)) and the source (***m***_*H*_(***x***)):

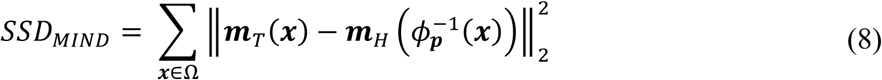

In the same fashion, for a multi-channel image, the original vector value of each pixel would be replaced by a matrix upon MIND transformation, and the dissimilarity of matrix-valued MIND representations would be expressed as the sum of squared elementwise differences. For a more detailed description on MIND see Heinrich et al [43].

### 2.5. Tensor Image Registration Library

Our novel image registration platform, the Tensor Image Registration Library (TIRL) uses the formalism laid out in *section 2.4* to aid fast prototyping of bespoke image registration pipelines for virtually any kind of images. All elements of the automated MRI-histology registration pipeline described in this paper were implemented in TIRL. The framework is based on Python 3.7, offering rich customisability via scripting. TIRL is a fully open-source project distributed as part of an upcoming release of the FMRIB Software Library (FSL) [53], with the intention to be used and further extended by community members. The main features of the library are summarised below.

TIRL follows an object-oriented programming paradigm. The registration process is realised by the interaction of several objects (Figure 4): the source and the target images (*TImage*), their domains (*Domain*), the coordinate transformations (*Transformation*), the cost function (*Cost*), the regularisation term (*Regulariser*), and the optimisation algorithm (*Optimiser*). TIRL defines custom file formats for saving any of the *TImage* (.timg), *Domain* (.dom), *Transformation* (.tx) and *TransformationGroup* (.txg) objects, which can later be loaded back into any compatible pipeline or used to extract quantitative transformation details after the registration is complete.

**Figure 4.**
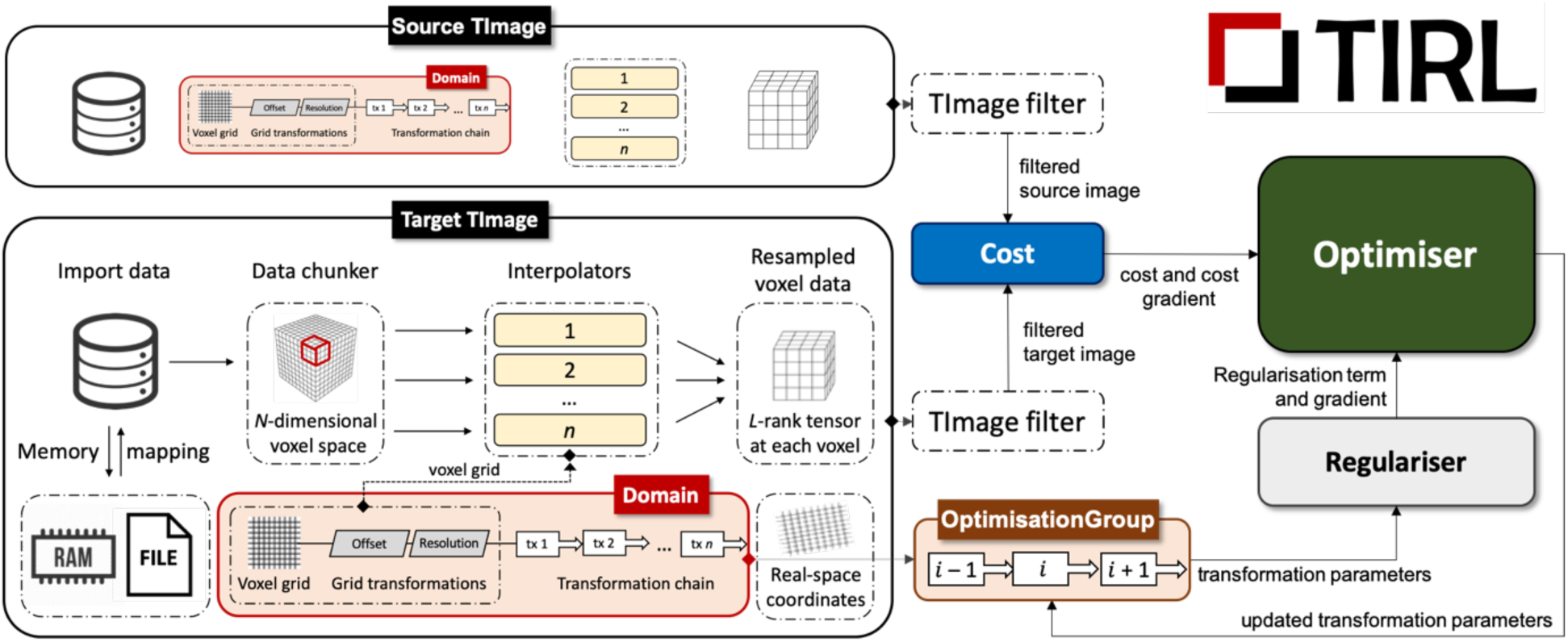
Schematic of the image registration logic within TIRL. Shapes with colour correspond to objects in the TIRL namespace. The source and target *TImage* objects comprise a memory-mapped container for raw (high-resolution) image data, a *Domain* object that defines the physical extent of the image by a chain of *Transformation*s, and an array of *Interpolator*s that map the high-resolution image data onto the current *Domain*. Both the target and the source images are passed through an image filter (to select a colour channel or calculate MIND *etc.*). The *Cost* object evaluates the filtered source image on the *Domain* of the filtered target image, and computes both the scalar cost and the cost gradient according to the object’s internal routine that is specific to the type of the *Cost* object (e.g. NMI, CR, SSD, *etc*.). These are fed into the *Optimiser* together with the regularisation term and regularisation gradients. The latter two are computed from the transformation parameters by the *Regulariser* object. The transformation parameters are pooled from the transformations of the target image chain that are selected by the *OptimisationGroup* object for simultaneous optimisation. (This object also allows transformations to be simultaneously optimised in both images.) Transformation parameters are updated in-place by the *Optimiser* object via the *OptimisationGroup*, until one of the convergence criteria is met (which is a parameter of the *Optimiser)*.

Similarly to the NIfTI standard for MR images [54], TIRL’s *TImage* is defined in physical space. This means that the image data is stored in a regular voxel grid with arbitrary number of dimensions (*N*), but each point in this voxel grid has an associate location in physical space (sometimes referred to as ‘millimetre-coordinates’). This is equivalent to the statement that each *TImage* is defined on a *Domain*. While NIfTI limits the mapping between voxel space and physical space coordinates to a single rigid/affine transformation (via the *q-form* and *s-form* matrices), the *Domain* object performs this mapping via a chain of *Transformation* objects, some of which may also be non-linear transformations. Hence, the position of a *TImage* in physical space can be manipulated by adding or removing *Transformation*s from *TImage*’s *Domain*, without changing the image data. Alternatively, the resolution of the image can be changed by evaluating the *TImage* on a new *Domain* that has the same physical extent but different matrix size.

A major advantage of the transformation chain formalism is that for example an affine transformation can be parametrised as permutations of scaling, rotation, shear, and translation transformations, and any of those components can be optimised separately and repeatedly at any point of the pipeline using the same syntax. Furthermore, the interpretation of transformation chains follows geometric intuition as opposed to affine matrices, especially in three dimensions.

A rich set of linear and non-linear transformations are currently implemented in TIRL. In particular, 3D rotations can be parametrised using *Euler* angles, rotation matrix, axis-angle, and quaternion formalism, with supported conversion between any two of these. The currently implemented options for non-linear transformations are polynomial coordinate transformations, and densely or sparsely defined displacement fields. This repertoire may be further extended by subclassing any of the *Transformation* objects.

The current version of TIRL implements the SSD, NMI, CR and *SSD*_*MIND*_ cost functions, as well as regularisation terms based on L1 and L2 norms, and membrane energy. TIRL is compatible with all gradient-free and gradient-based optimisation algorithms available from the SciPy [55] and NLOpt [56] optimisation libraries. Custom implementations of cost functions, regularisation terms, and optimisation algorithms are also supported via the *Cost, Regulariser*, and *Optimiser* base classes.

Image masks are often used in neuroimaging on diseased or artefacted images to downweight the cost for these regions, preventing erroneous registration. In TIRL, masks can be specified for each *TImage*, and these are adaptively combined during registration to weight the cost function for the intersection of the target and the source image.

TIRL supports easy importing from various file formats and provides an all-compatible workflow for various kinds of images (e.g. scalar-, vector-, matrix-, tensor-valued images with arbitrary number of dimensions) via the *TImage* object. It is an *N*-dimensional image, in which every voxel value is an *L*-rank tensor. For large images that do not fit in memory, the data of the *TImage* is dynamically retrieved from a memory-mapped binary file that resides on the hard drive. Furthermore, all *TImage* operations (including interpolation) are automatically chunked and parallelised for faster computation.

We implemented all three steps of the proposed MRI–histology registration pipeline in TIRL because it enables fast prototyping and rich customisation of bespoke image registration pipelines and has the flexibility to work with mixed sets of 2D and 3D images.

### 2.6. Stage 1: Histology image to tissue block photograph

The registration between a histological image and the corresponding tissue block photograph can be formalised in either direction. Morphing the domain of the histology image to the tissue block has the advantage of providing forward mapping towards the MRI end of the pipeline, hence, it is not necessary to invert any transformations to find a certain histological feature in MRI space. On the other hand, resampling the histology image and the MRI image on any of the photographic intermediates requires less non-linear deformation from both ends, resulting in a more symmetric registration approach with potentially less inverse-consistency error [57]. We therefore adopted the latter formalism, and denoted the block as the target, and the histology image as the source.

In line with Equation (1), we define the inverse transform (the one that maps the target domain onto the source domain) as a chain of *Transformation* objects acting on the *Domain* of the target *TImage* (tissue block photo). As shown by the top bar in Figure 5, the chain comprises 2D scaling, 2D rotation, 2D affine, 2D translation, and 2D deformation. The chain is prepended with a *Translation* object that moves the pixel at the centre of the image to the origin, ensuring that the first chain operation is applied on centralised pixel coordinates. The order of *Transformation* objects within the chain follows the intuition behind aligning the images by hand. Rotations come after scaling to ensure that the image is stretched along the original pixel axes, and rotations precede translations to ensure that rotations are carried out about the centre of the image, not some arbitrary centre of rotation. In TIRL, it is possible to simultaneously optimise any subset of transformations in the chain. By optimising transformations in the order of increasing degrees of freedom (DOF), previously optimised coarser transformations provide a more suitable initialisation for finer transformations that are optimised later. This increases the chance of finding the global optimum by local optimisation methods, which are generally faster than global optimisation methods. When choosing transformations for simultaneous optimisation, it is important to avoid optimising redundant parameter pairs against each other, such as the components of a full affine matrix against rotation angles, as this would create infinitely many equal minima in the cost function and lead to undetermined behaviour of the optimiser. In this particular case, rotation angles would have to be optimised first to achieve a coarse initialisation, and held constant while the components of the affine transformation matrix are fine-tuned, accounting for both shears and finer rotations. Based on these considerations, stage 1 optimises the above transformation chain in four steps: (1) rotation search, (2) rigid registration with anisotropic scaling (“pseudo-rigid registration”), (3) affine registration, and (4) non-linear registration. Now we discuss these optimisation stages in greater detail.

**Figure 5.**
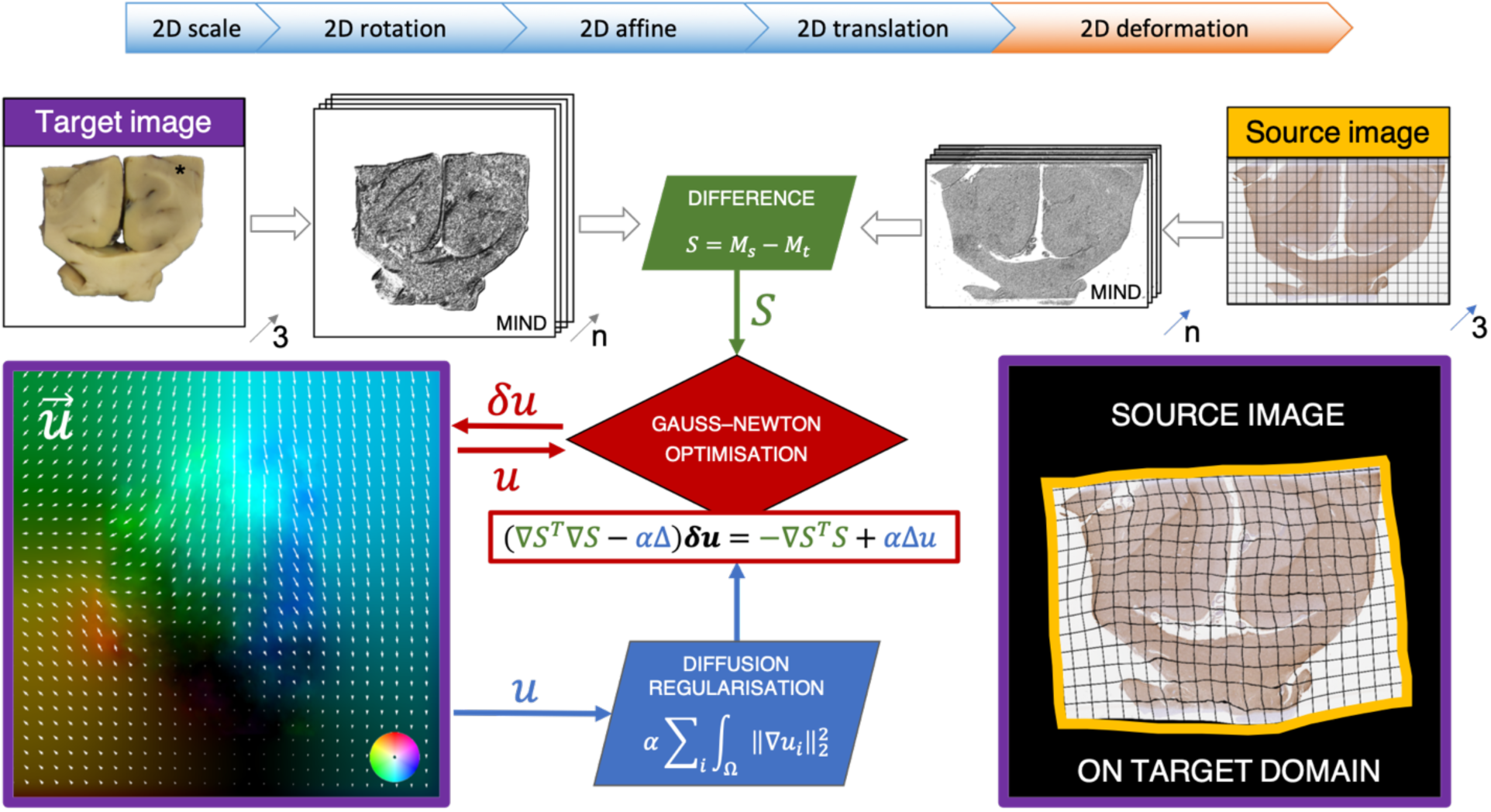
Overview of stage 1: histology-to-block registration. *Top bar*: transformation chain of the target image domain. The chain is optimised in four steps: rotation search (2D rotation), rigid registration with anisotropic scaling (2D scale, 2D rotation, 2D translation), affine registration (2D affine), and non-linear registration (2D deformation). *Flowchart*: non-linear registration of a histological image to a tissue block photograph. The target and source RGB images are flattened to grayscale during pre-processing, and MIND-filtered, yielding images with n=8 channels. An 8-channel difference image is calculated by resampling the filtered source on the target domain and subtracting the filtered target. The non-linear transformation is parametrised as a dense array of displacement vectors (***u***(***x***)) over the target domain and is initialised to zero for all ***x***. Since the displacement vectors are optimised independently, diffusion regularisation is applied to prevent the source image from folding over itself and leading to a non-diffeomorphic mapping. Local updates to the displacement vectors are computed at every iteration by the Gauss–Newton optimiser, based on the 8-channel difference image (and its derivative) and the regularisation gradient that are supplied by the *Cost* and *Regulariser* objects. As a result of the optimisation, the resampled histology image (source) becomes maximally similar to the block photograph (target).

(1) *Rotation search*: a line search with constant 10° increments is conducted around the full circle to find the best initial rotation for the images. In our experiments we found that 10° was a good compromise between computational performance and the robustness of the initialisation. The search maximises NMI using Powell’s method [58] at 0.5 mm isotropic resolution. (2) *“Pseudo-rigid” registration*: starting from the three best initial rotations, a 5-DOF registration (2D rotation + 2D translation + 2D anisotropic scaling) is carried out using *SSD*_*MIND*_ as the cost function, and the BOBYQA optimiser [59] at 0.5 mm isotropic resolution. Similar tests carried out by *Osechinskiy* et al [40] had suggested the use of the NEWUOA optimiser, but we found the convergence properties of its bounded variant, BOBYQA more reliable for this purpose. We also found that MIND-based rigid registration was robust enough to identify some major defects in the structural correspondence of the images (e.g. when a piece of tissue is torn at the corner during histological processing). To find these regions, the binarized source image was subtracted from the binary target and the identified regions were sorted by area. Anything larger than 5% of the block surface was added to the target mask as a zero-filled region to exclude it from the computation of the cost. (An example mask is given in the supplementary material.) In TIRL, the masks from the source and target image are combined and used as a multiplier for the cost and – in case of gradient-based optimisation – the cost derivative, desensitising the optimisation algorithm to changes in an area where the mask value is close to zero. The best rigid initialisation was chosen from the three results on the basis of minimum final *SSD*_*MIND*_ cost, and fed into *the affine registration step* (3) at 0.5 mm, 0.25 mm and 0.1 mm resolutions, optimised for *SSD*_*MIND*_ using the BOBYQA optimiser.

The flowchart in Figure 5 summarises the concluding step of stage 1: optimising *the non-linear transformation* (4). This step employs diffusion registration that was introduced by *Fischer* and *Modersitzki* [52, 60, 61] and further adapted for the *SSD*_*MIND*_ cost function by *Heinrich* et al [43]. Here we describe the optimisation process with a linear pre-alignment step, as it is implemented in TIRL. All equations below follow the denominator layout convention, and all vectors are column vectors unless otherwise stated.

The non-linear transformation is preceded by a sequence of linear transformations, which can be combined into a single affine transformation (**A**) that operates on homogeneous coordinates (***x***). (In TIRL, consecutive linear transformations are automatically replaced by the equivalent affine transformation to optimise the computation of new image coordinates). The non-linear transformation is parametrised as a deformation field ***u***(***x***), that incurs additional displacement to the linear mapping of target coordinates (***x*** ∈ Ω) to source coordinates (***x***’ ∈ Ψ):

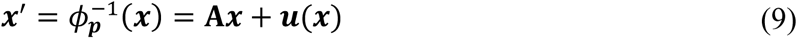

Given **A**, the optimisation aims to find plausible values for ***u***(***x***) that together with **A** minimise the overall difference between the MIND representation of the two images. Mathematically, this is equivalent to minimising the following cost functional:

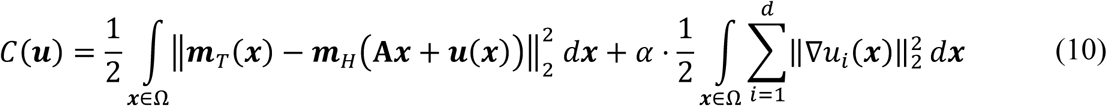

where the first term is the cumulative Euclidean distance of MIND vectors representing image dissimilarity within the target domain, and the second term is the so-called diffusion regularisation penalty that is supposed to prevent unrealistic folds in the transformed source image by penalising sharp gradients in each component (*i* = 1, …, *d*) of the deformation field. The *α* parameter is a weighting factor that controls the relative importance of the regularisation with respect to image dissimilarity.

The optimisation proceeds by iteratively computing vector updates (***h***(***x***)) of the deformation field until the desired precision (*ϵ* = 10% of the pixel size) or the maximum number of iterations (*k*_max_ = 20) is reached:

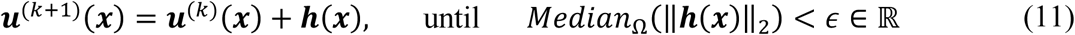

Using the discretisation in (11) we now aim to reformulate (10) such that it becomes linear with respect to the updates. We therefore rewrite the argument of the dissimilarity term in (10) using (11) and first-order Taylor expansion:

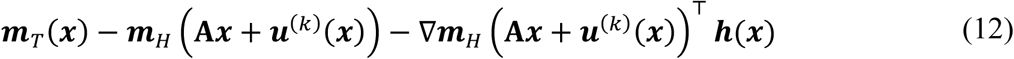

As deformations grow larger, the linear approximation based on the originally computed MIND vectors of the source image (***m***_*H*_) becomes less accurate. To avoid this, we recompute the respective MIND vectors at every iteration from the transformed source image, for which we introduce the 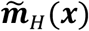 notation:

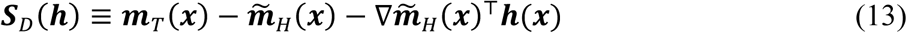

The regularisation term is already linear with respect to ***h***(***x***), because the differential operator 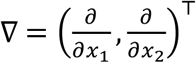 is linear. Substituting (11) into the regularisation term of (10) yields:

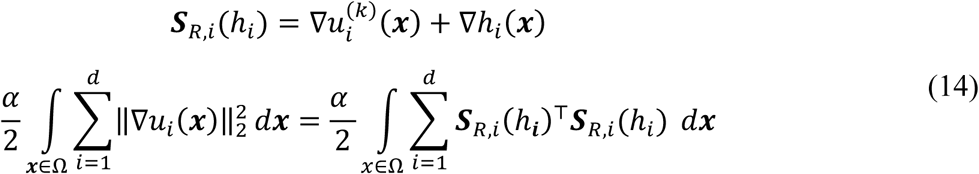

After substituting (13) and (14) into (10) we obtain the following linearised expression for the cost at the *k*-th iteration:

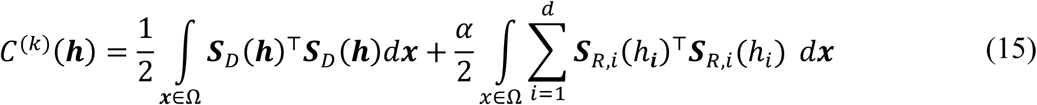

To minimise the cost of the *k*-th iteration (15), we formulate the corresponding system of *Euler–Lagrange* equations for each spatial component of the deformation field (*i* = 1, …, *d*):

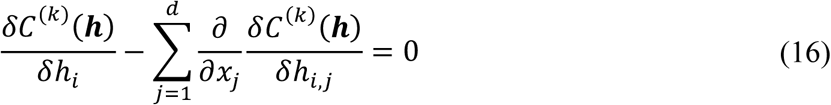

Expressing the functional derivatives in (16) yields the following system of equations (*i* = 1, …, *d*) for each pixel in the target domain:

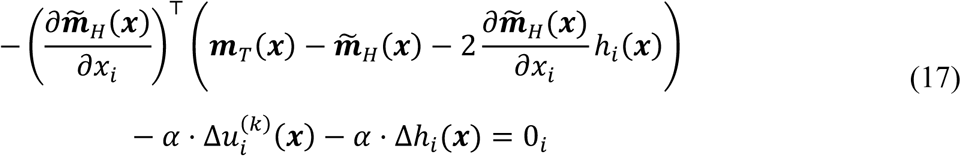

where Δ is the Laplace operator (discretised using a 4-point stencil on the 2D target grid). By rearranging (17), substituting ***S***_*D*_ from (13), and introducing the notation 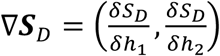 for the *Gateaux*-derivative, we obtain the following formula, which is equivalent to minimising the cost functional in (15) by the *Gauss–Newton* method [62], where ***h***(***x***) constitutes the update step:

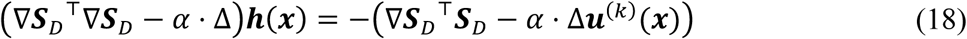

Equation (18) is eventually solved simultaneously for all pixels in the target domain for ***h***(***x***) using the sparse solver of SciPy [63]. After every iteration, the deformation field is updated according to Equation (11). At the end of the optimisation the estimated deformation field is used to initialise a new optimisation at a higher grid resolution and so forth. In our experiments, sufficient convergence was reached in 20 iterations at each of the 1.5 mm, 1 mm, 0.5 mm and 0.25 mm resolution levels.

To test the accuracy of this stage, we registered H&E, ferritin- and PLP-stained histological images to 6 corresponding tissue blocks from various anatomical locations. We selected all available stains that exhibited visible grey-white matter contrast. The blocks were selected to represent the observed variability of the size, shape, and anatomical texture of the blocks in the full dataset. The pre-processed images were imported to TIRL, and the grayscale histological images were thresholded between 150 and 400 to remove shadows on the slide edge (inherent to slide scanning) and the white background. The histology image was further smoothed by a Gaussian kernel (*σ* = 0.1 mm) partly because voxel-to-voxel variations due to staining cell nuclei are not represented in the block photo, and partly to prevent small holes arising from the thresholding creating a false texture that MIND is sensitive to. The resolution of the images was set to the resolution of the photograph, and image centres were moved to the origin. Before computing the MIND representation, the images were flattened to a single channel as described earlier.

The registrations were evaluated in terms of maximum and average registration error of overlapping contours. Grey-white matter contours were defined separately for the histology images and the block photos by manually annotating approximately 200 points for each image in Fiji [64]. Pairwise alignment of the contours was assessed after registering the images. To establish pairwise point correspondence between the contours, point coordinates were parametrised, and all contours were upsampled by B-spline interpolation to comprise exactly 2000 points. The contours from the block photographs (target) were transformed into the domain of the corresponding histology images (source) by the transformation chain of the target images. In the histology domain two parameters were computed for every pair of contours: (1) the *Hausdorff distance*, and (2) the *median contour distance*. The former measures the maximum distance between corresponding contour points with the same index, yielding an estimate of the largest registration error. The latter is calculated by measuring the distance of each contour point from the closest point of the other contour and taking the median of these measurements. As this measure is independent from point correspondence errors that arise from the manual nature of the segmentations, it provides a more realistic quantitative estimate of the overall registration error, which can be qualitatively observed by eye.

### 2.7. Stage 2: Tissue block to brain slice photograph

The second step of the automated registration pipeline maps the coordinates of the tissue block photograph (Figure 6F) to the domain of the brain slice photograph (Figure 6A). Given that the tissue block is significantly smaller than the whole-brain coronal slice, it is computationally more efficient to choose the tissue block as target, and the brain slice as source, as the latter will be repeatedly interpolated at the target domain as the optimisation proceeds. The vastly different size of the objects however also poses a registration challenge, as extreme oversizing or shrinkage of the block image can be a trivial solution to minimise SSD cost based on pixel intensities. Furthermore, most brain slice photos exhibit high degrees of self-similarity: a block with a certain anatomical pattern may be a relatively good fit at multiple positions. Without prior initialisation, this would require extensive spatial search or a very time-consuming global optimisation method to succeed, especially when multiple blocks must be registered to the same slice.

**Figure 6.**
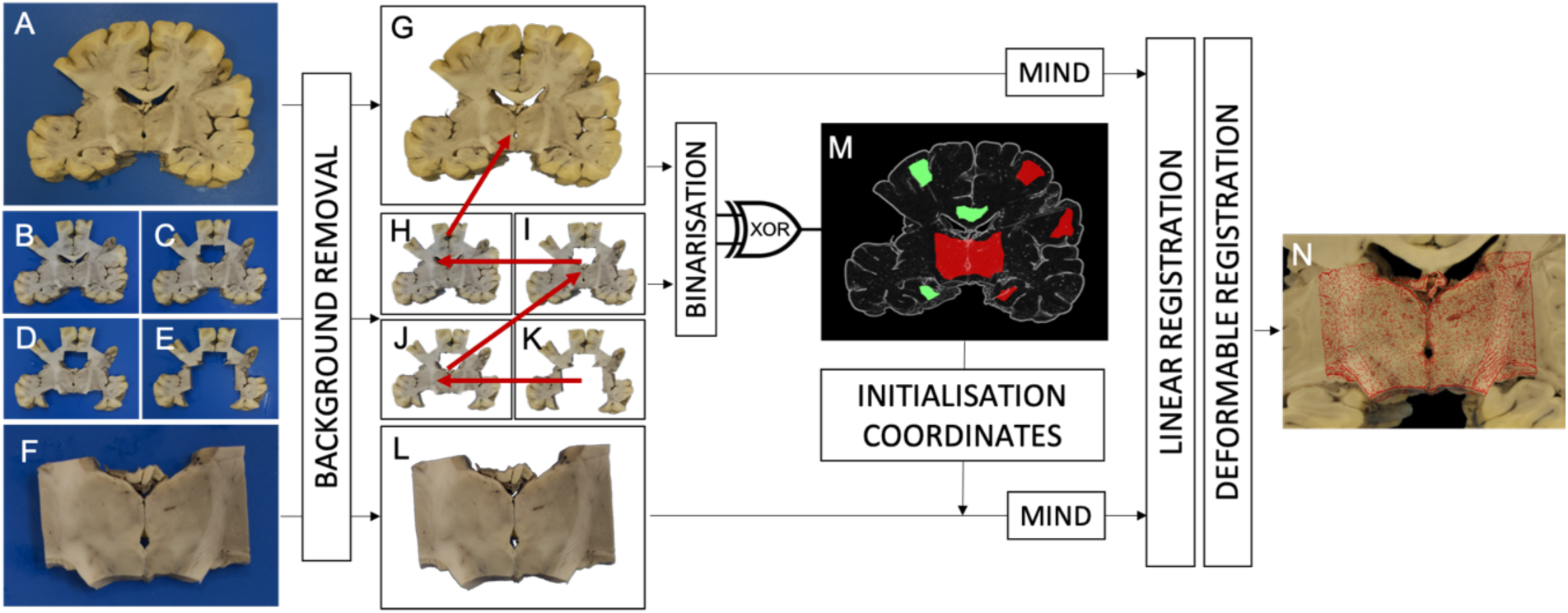
Overview of stage 2: block-to-slice registration. Registration of a coronal slice (A) to a tissue block (F). Auxiliary slice photographs (B-E) were taken every time that a new tissue block was sampled from the same coronal brain slice. The blue background was removed from the photographs using *k*-means classification of RBG vectors (*k*=2). The segmented coronal slice (G) served as a common reference domain for affine registering all segmented auxiliary slices (H-K). Affine registered auxiliary slices were pairwise non-linearly registered in reverse order, starting with the most damaged slice (red arrows). The original coronal slice and the aligned auxiliary slices were binarized, and the XOR operation was applied to each consecutive pair of them to identify non-matching areas as insertion sites (M). Following rigid initialisation at the centroid of each insertion site, the background-segmented block photograph (L) was affine registered at each site. At the site where the affine registration yielded minimum *SSD*_*MIND*_ cost, a non-linear deformation was also performed to achieve accurate alignment of the block with the intact brain slice photo (N).

To keep stage 2 automated and relatively fast, prior information is obtained from a series of autopsy photographs (“cut-out images” or “auxiliary slices”, Figure 6B-E) capturing the original brain slice (Figure 6A) after the excision of a tissue block (or multiple spatially-distinct blocks). These photographs are labelled in chronological order of the dissection process. Each cut-out image is affine registered to the original coronal image first, then consecutive pairs of the cut-out images are registered by a chain of affine and non-linear transformations. The registered images are binarized and the binary difference (XOR) is taken to identify possible “insertion sites” for the blocks (Figure 6M).

The registration of the consecutive slices follows the same scheme as in stage 1: with the former cut-out image as target, and the later image as source, the algorithm performs an initial rotation search, followed by pseudo-rigid, affine, and non-linear registrations. For the rotation search, we used NMI cost as it is computationally less demanding than MIND, and it proved to be robust enough for the purpose. For the rest of the process, we used *SSD*_*MIND*_, which is more sensitive to texture. Furthermore, the scale parameters of the “pseudo-rigid” step were confined to the range 0.9 – 1.1 and the BOBYQA bounded optimisation algorithm was used to prevent the oversizing/shrinking of blocks with less salient anatomical pattern. To identify insertion sites, each of the registered images were binarized by clipping intensity values at 1, and the images were multiplied to create a segmentation of non-aligned parts. The segmentation result was eroded by 5 mm, and the centroid of all connected components above an area of 1 cm^2^ were denoted insertion sites.

To register multiple blocks automatically to the same slice, the search for insertion sites was performed once, then each block was affine registered at each site. The affine registration that yielded minimum *SSD*_*MIND*_ cost at the end of this stage was used to initialise a non-linear registration step for further refinement, concluding stage 2 for each block.

### 2.8. Stage 3: Brain slice photograph to MRI volume

In the final step of the pipeline, the 2D coordinates of the brain slice photograph are mapped into 3D MRI space. The choice of the brain slice as target and the MR image as source is obligatory, as the interpolation of a 2D image on a 3D domain would not only be extremely inefficient to perform at each iteration of the optimisation process, but is also ill-defined. Stage 3 instead makes gradual improvements to the position, orientation, in-plane deformation and out-of-plane curvature of the brain slice photograph and compares it with the MRI values that are resampled from the intersection of this warped 2D domain with the MRI volume. The four steps of stage 3 and the respective transformation chain are illustrated in Figure 7. These transformations were optimised in various combinations with gradually converging boundary constraints, which are described below for each step.

**Figure 7.**
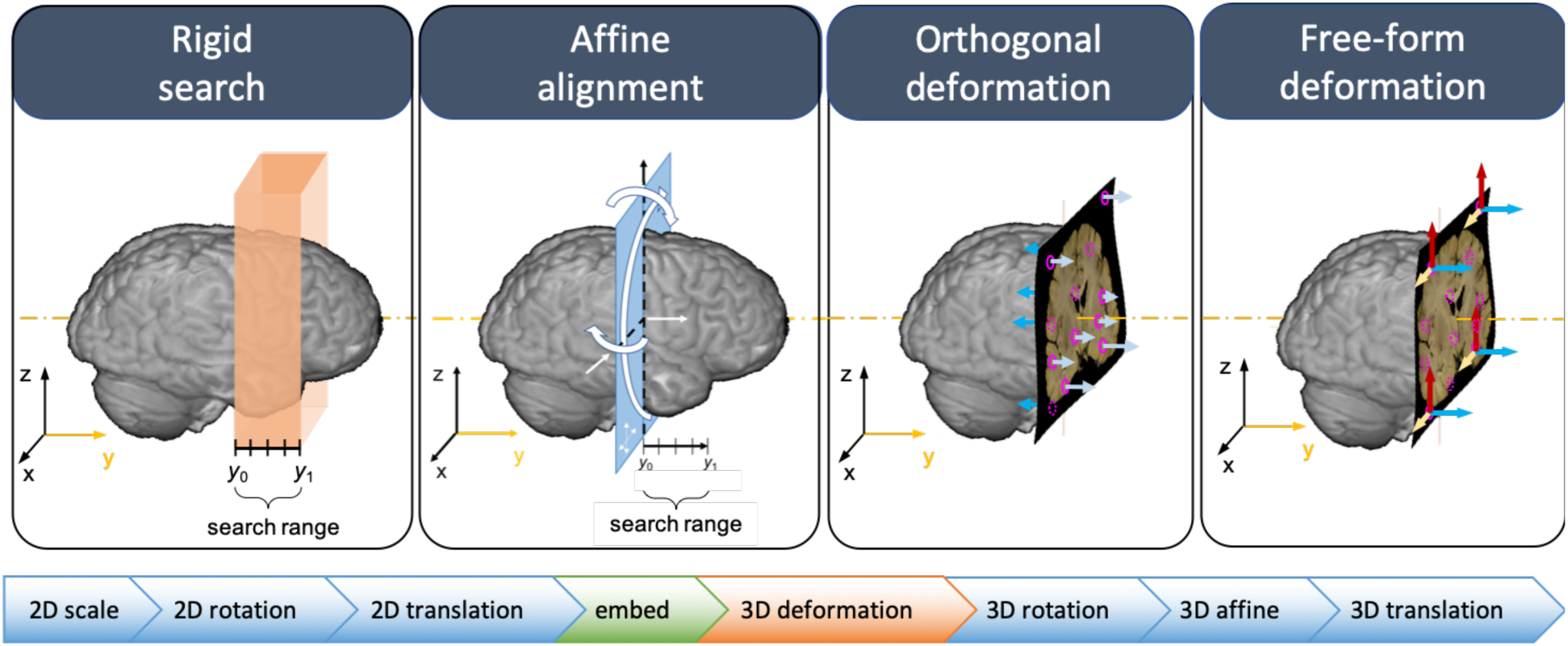
Overview of stage 3: slice-to-MRI registration. *Panels from left to right*: Consecutive steps of optimising different subsets of transformations in the transformation chain (*bottom bar*) of the brain slice photograph. *Rigid search*: a rectangular search volume is defined in MRI space by its centre, orientation and thickness (*orange slab*). The slice photograph is repeatedly initialised at various positions along the central axis of the slab, while the best 8-DOF (3D rigid + 2D scaling) alignment is sought. As the slices are coronal, their *z*-axis is initially aligned with the *y*-axis of the MRI. *Affine alignment*: 3D affine parameters are optimised to refine the linear registration. *Orthogonal deformation:* 50 control points (*pink*) are defined on the domain of the slice photograph. By dynamically changing the local protrusion/retraction of the slice at these points, the slice is slightly curved to compensate for off-plane distortions. Membrane energy is employed as a regularisation penalty to prevent sharp bending. *Free-form deformation*: the same 50 points are allowed to move freely in 3D space to compensate for in-plane deformations. (Arrows are shown only for a subset of the control points for better visibility.) In the last two steps, the displacements of the control points are constrained by membrane energy.

Step 1 aims to find the best rigid alignment of the brain slice image within a confined, approximately 2-cm thick rectangular search region in MRI space by iterating a 2D and a 3D bounded optimisation of *SSD*_*MIND*_ cost (using BOBYQA), namely Step 1A (2D scaling, 2D rotation, 2D translation) and Step 1B (2D scaling, 3D rotation, 3D translation). Both step 1A and step 1B are carried out twice at each resolution level (4 mm, 2 mm, 1 mm, 0.5 mm): first with slight Gaussian smoothing (*σ* = 1 px), next without smoothing: [1A+1B]_2_ The search region is most conveniently defined manually by its centre (3D coordinates), orientation (3D vector) and thickness (scalar), seven parameters altogether. In our case the standardised dissection strategy allows the definition of the slab automatically by slice number, given that all slices are coronal, separated by 1 cm along the anterior-posterior (A-P) axis of the brain, starting from the plane of the mamillary bodies toward the anterior and the posterior poles of the brain. The normal vector of the slab is collinear with the A-P axis, unless the brain is significantly tilted. Stage 1A and 1B are then performed with the same initial parameters at 5 equally spaced points along the short axis of the search range, and the position with the least *SSD*_*MIND*_ cost at the end is accepted as the best initialisation position for the slice. At this location, stage 1A and 1B is repeated for a final time, but the optimiser is changed from BOBYQA to its closest unconstrained equivalent, NEWUOA. In our experience, the rotation and scale parameters have a stronger influence on the cost function, therefore translation parameters are effectively only optimised late in the process after these two. Unconstrained optimisation by NEWUOA with the nearly optimal rotation and scale parameters allows escaping the current slab position that might have been reached with these parameters being suboptimal in the previous iterations. The result of this unconstrained optimisation is accepted as the best pseudo-rigid alignment of the slice photograph, which is then fed into step 2.

Step 2 aims to compensate for shears of the slice photograph and optimises the parameters of the 3D affine transformation. Given the pseudo-rigid alignment from step 1, the scale, rotation, and translation parameters are not expected to change substantially in this step, and strict bounds are set for these parameters in the BOBYQA optimiser.

Step 3 introduces deformations that are orthogonal to the slice photograph to account for the slightly irregular, non-planar nature of free-hand cuts through the brain. The assumption behind step 3 is that variations in the anatomical pattern of neighbouring slices due to off-plane deformation is a larger contributor to the misalignment after affine registration than in-plane deformations are, as long as the resolution is coarse (4 mm, 2 mm). In-plane deformations are therefore not introduced until step 4. The transformation of step 3 is parametrised by orthogonal (*z*-axis) displacements at 16 ≤ *N*_*c*_ ≤ 128 control points (nodes) defined on the domain of the 2D brain slice photograph (Figure 7, panel 3). The number of points was determined empirically as a trade-off between registration accuracy and computation time. The displacements for the rest of the image domain are calculated from the known *z*-axis displacements using interpolation by a set of Gaussian radial basis functions (*G*(*r, σ*)):

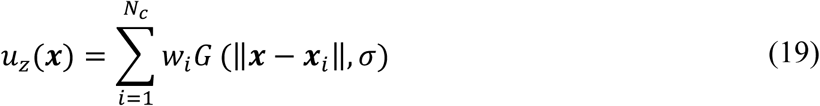

The *σ* parameter defines the effective radius of each node, and is set to the average Euclidean distance between all pairs of control points. The weight for each node is calculated by fitting the above equation for the predefined *z*-axis displacements at the control points. The parameters (*z*-axis displacements) of the transformation are optimised for minimum *SSD*_*MIND*_ cost using the NEWUOA algorithm. In addition to the *SSD*_*MIND*_ cost, a membrane energy regularisation term was included in the cost function to prevent sharp bends of the domain. Membrane energy was defined at any point ***x*** of the target domain as the L2-norm of the second derivatives of the local deformation field, and summed over the entire domain to obtain the full regularisation term:

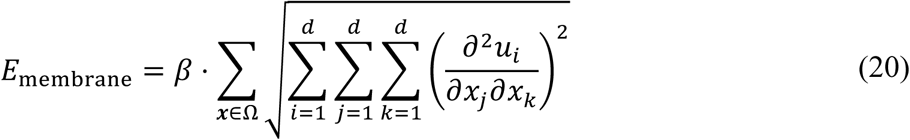

where *β* is a regularisation parameter that was set to 10^−8^.

In step 4, we extend the transformation from the previous step such that deformations at each point of the target domain are simultaneously optimised along all three spatial dimensions. In addition to the in-plane deformations that naturally occur during handling the brain, through-plane deformations of step 3 also incur in-plane deformations by projecting 3D MRI data onto a regular 2D grid. The tangential deformation components (*u*_*x*_, *u*_*y*_) are initialised to zero and calculated for all pixels of the target domain from the respective deformation components at the control points throughout the optimisation. Previously optimised orthogonal deformations (*u*_*z*_) are retained, but may also change in step 4, rendering this final transformation a free-form deformation with restricted degrees of freedom that is dictated by the number of control points. Membrane energy regularisation was also used at this stage.

While the control points can be defined manually, to keep the pipeline automated, we generated quasi-random two-dimensional coordinates for the control points by drawing numbers from a sequence of rational fractions (Halton sequence [65]). The advantage of using Halton-sequence points versus a rectangular grid is that the points provide similar uniform coverage of the area, while the number of points can be set to any positive integer. This is important, because the complexity of the optimisation increases with the number of parameters, and for the same fine-grain control over the deformations, one would need more points in a regular grid layout than with Halton points. When compared to pseudorandom point placement, Halton points have the advantage of being deterministic (reproducible) and having low discrepancy (no two points are extremely close to each other). To restrict deformations to the area of the coronal brain slice (excluding most of the background), a bounding box was defined for the brain slice by Otsu-thresholding and it was expanded by 10% in each direction. The Halton points were finally scaled and shifted accordingly to fit inside the bounding box.

### 2.9. Combining transformations

Given all three previously described stages of the pipeline, there are two competing alternatives for combining them to achieve end-to-end MRI-histology registration. While these methods are equivalent in theory, they are slightly different in terms of their implementation in TIRL and their practical consequences.

The first method is forward mapping by concatenating the optimised transformation chains of the target images from each step (histology, block, slice). The practical consequence of this method is that the histology data is never interpolated (unless subsampled to a lower resolution at the start), and that histological coordinates can be mapped to MRI coordinates without the necessity to invert any of the optimised transformations. However, the deformations between MRI and the histology image can be large and may not be estimated accurately in every case by the combination of multiple non-linear transformations.

In the second method, instead of concatenating non-linear transformations, the registration converges at the half-way point, on the tissue block photograph. The optimised transformation chains from the second and third stages are concatenated, and the MRI data is evaluated on the domain of the block photograph by interpolation. The difference from the first method is that the histology data must also be evaluated on this domain by interpolation using the optimised transformation chain from the first stage. The main advantage of this method is that the non-linear transformations are not chained, consequently the registration error incurred by each of them is not amplified by the other, which means that the final alignment can be more accurate. The practical consequence of this method is that the original histological coordinates cannot be mapped to MRI space without inverting a non-linear transformation, the precision of which may be affected by the inverse consistency error [57].

For the practical purpose of correlating MRI and histology parameters in a predefined region of interest, the limitation of the second method is not relevant as long as pixels from the images can be overlaid at any desired resolution. We therefore adopted the second method to achieve greater precision in estimating tissue deformations.

## 3. Results

### 3.1. Stage 1 results: Histology image to tissue block photograph

The accuracy and robustness of stage 1 was tested on 6 tissue blocks from various anatomical regions of the same human brain: (1) the orbitofrontal cortex, (2) the anterior cingulate cortex, (3) the anterior limb of the internal capsule (also including parts from the caudate nucleus and the putamen), (4) the hippocampus, (5) the thalamus, and (6) the visual cortex at the banks of the calcarine fissure. Each of the blocks had corresponding histological sections stained with H&E, for ferritin and for PLP. (1) and (6) were further stained with LFB+PAS. The rest of the immunohistochemistry images were not considered for registration with the tissue block photograph, as they exhibited virtually no grey-white matter contrast. Registrations also were performed with various regularisation weights (0.2 – 2.0) to test the effect of regularisation on registration accuracy.

Figure 8A-D shows a representative example how the rigid, affine and non-linear transformations gradually improved the registration accuracy between the histological image and the tissue block photo during stage-1 registration. To gauge the plausibility of the non-linear transformations associated with different regularisation weights, the Jacobian determinant maps were plotted in Figure 8E-F for a small (*α* = 0.4) and a larger (*α* = 1.2) regularisation weight. At each pixel, the Jacobian determinant describes the local shrinkage/dilation of the image elements by the non-linear transformation. The range of the Jacobian values was 0.1 – 2.5 for *α* = 0.4 and 0.8 – 1.2 for *α* = 1.2. As none of these values were below zero, the diffeomorphic nature of the transformation was preserved: no image elements were lost by abnormal self-folding of the image in either case. However, in the under-regularised case, the sharp transitions between large local deformations are most likely driven by local intensity variations in the images, pointing at the non-physical nature of this transformation. On the contrary, in the more regularised case the deformations were smaller, more balanced and more homogeneous, reflecting actual tissue deformations. This effect is further evidenced by the violin plot in Figure 8G, which shows the distribution of deformations (in millimetres) versus regularisation weight. Irrespective of the regularisation weight, the deformations were roughly evenly distributed between 0 and 1 mm, which seems a physically plausible estimate for the magnitude of tissue deformations (the deformations of the background were excluded). Increasing the regularisation weight restricted large local deformations, which is seen as a reduction of the heavy tail in the plots of Figure 8G.

**Figure 8.**
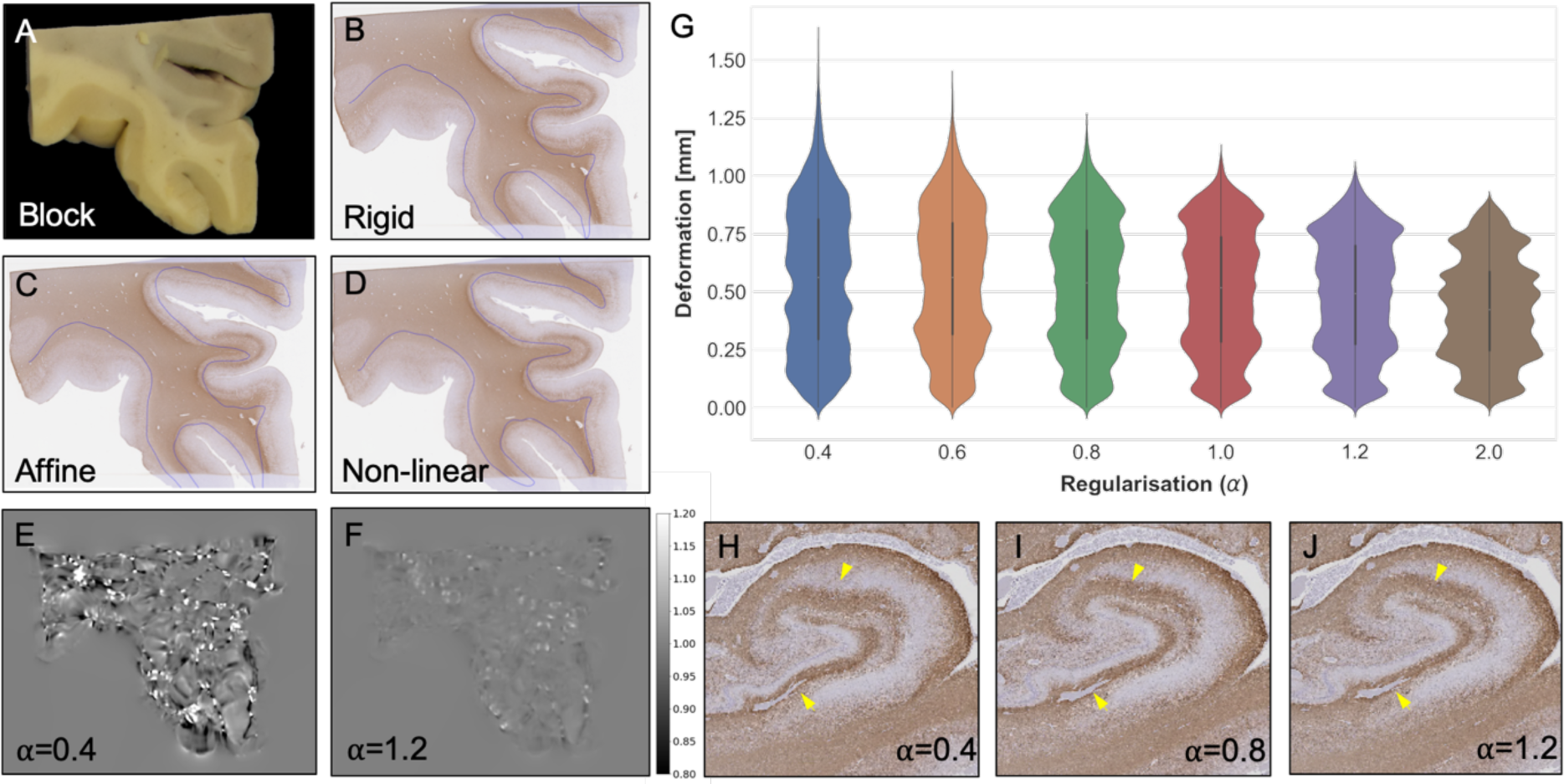
Results of stage 1 (histology-to-block registration). *(A)* Photograph of the posterior surface of a tissue block from the left orbitofrontal cortex (OFC) that was used as the target image for the registration of the corresponding histological image. *(B-D)* PLP-stained histology image of the same region shown with the transformed overlay (*blue curve*) of the manually defined grey-white matter contour of the block photograph after each registration step. The registration accuracy is gradually improved by each consecutive step of the optimisation (pseudo-rigid, affine, non-linear) as evidenced by better alignment between the overlay and the grey-white matter boundary of the histological image. *(E)* The Jacobian map of the non-linear transformation over the OFC indicates large (beyond 20%) local shrinkages and dilations for *α* = 0.4. *(F)* The Jacobian map for *α* = 1.0 over the OFC shows physically plausible shrinkage and dilation (both ∼20%) of the tissue. *(G)* Typical in-plane deformations in millimetres as a function of regularisation weight for the registration shown in A-D. Increasing the regularisation weights restricts implausible large local deformations that is seen as a reduction of the upper tail. *(H-J)* Distortions shown on the PLP-stained histology image of the hippocampus as a function of the regularisation weight (*α*). Too little regularisation (*α* = 0.4) yields jagged appearance of the anatomical contours after registration (*yellow arrowheads*), whereas regularisation weights above 0.8 yield physically plausible, almost identical results.

Under-regularised stage-1 registration distorts anatomical contours (Figure 8H), which is hard to notice unless compared against the result of well-regularised registrations (Figure 8I-J). Minor inaccuracies like this are almost undetectable by eye or even by comparing grey-white matter contours. Nevertheless, the deformation field carries an additional rotation component (curl ***u***(***x***)) around the distorted regions. This may locally bias the histology-derived fibre orientations and insidiously reduce the correlation with MRI-derived fibre orientations even when the registration appears grossly accurate. For direction-sensitive applications it is therefore recommended to set *α* as high as reasonably possible, even at the expense of a slightly higher overall registration error. In our experiments, we observed no obvious anatomical distortions for *α* > 0.8.

Figure 9 shows the accuracy of stage-1 registration in terms of the two distance measures for each histological stain, for each optimisation step (rigid, affine, non-linear), and for multiple regularisation weights (0.2–2.0). *Hausdorff*-distances were generally larger (1.0–2.5 mm) than the perceived registration error, which was more accurately captured by the median contour distance (0.2–0.7 mm). The reason is that the former strongly depends on accurate point correspondence between the contours, which cannot be guaranteed given the irregularities of hand-segmentation and the resultant difference in the length of the contours.

**Figure 9.**
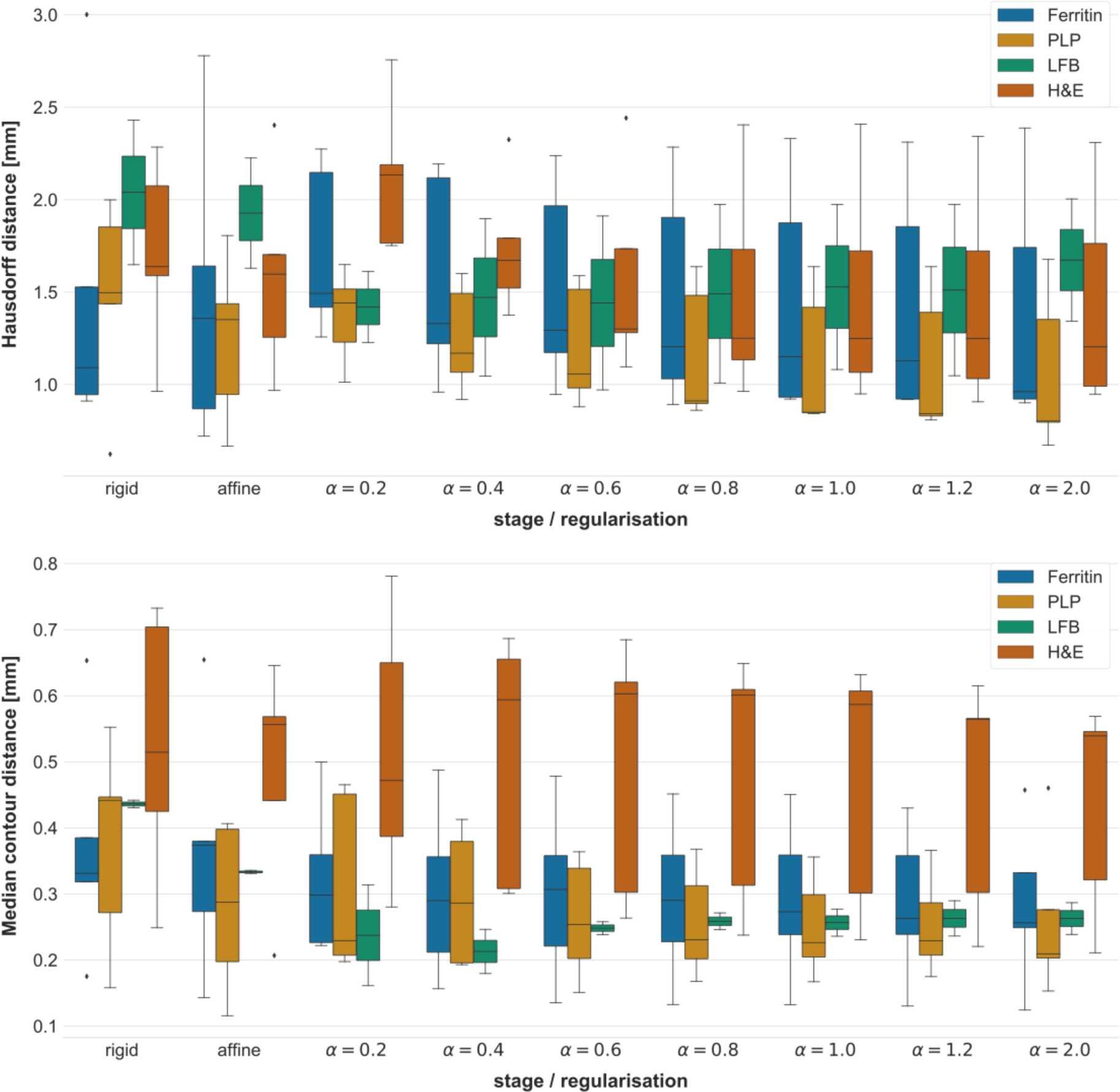
Accuracy of stage 1 (histology-to-block registration). The boxplots show the *Hausdorff*-distance (*top*) and the median contour distance (*bottom*) between the grey-white matter contours of the registered images. Both distances are reported in millimetres for each step of the registration (including multiple regularisation weights for the non-linear step) and for each histological stain (*n*=6, except for LFB *n*=2). Indicated on the bars are the median, the interquartile range and the extrema of these measures. The Hausdorff-distance is a biased estimator of the largest registration error (cf. text in *section 3.1*), whereas the median contour distance is a more accurate representation of the visually perceived accuracy of the registration. Based on the latter measure, the most consistent registration results could be obtained with the LFB stain, the most accurate results with the PLP stain, and the most inaccurate results with the H&E stain.

As evidenced by median contour distances, the rigid, affine and non-linear steps of the registration gradually improved the alignment of the tissue block photo with the PLP, LFB and ferritin-stained histological images, and the accuracy of the registration was similar for all three stains (0.2–0.3 mm). However, the same improvement was not seen with the H&E sections, for which the best results were achieved with the rigid alignment (0.4–0.7 mm). This is most likely explained by the grey-white matter contrast that was high with the former three stains, and almost absent in H&E stained sections. Based on these results, successful stage-1 registration requires at least one stained section for each region of interest that has comparable contrast properties to the MRI image. The rest of the histological sections can then be registered linearly to this section.

Based on the median contour distances, the most consistent results could be achieved with LFB+PAS-stained sections. In the physically plausible regularisation range (*α* > 0.8) the accuracy was consistently 0.25–0.28 mm. Slightly more accurate registrations could be achieved with the PLP stained sections, where the best results (0.20–0.28 mm) were obtained with *α* = 2.0 regularisation. The best results with the ferritin-stained sections (0.25–0.34 mm) were also obtained using *α* = 2.0 regularisation.

While running stage 1 registration, we encountered a few unexpected results. Most notably, the sample from the anterior cingulate was too large for a standard histological slide (25 x 75 mm), and the superior portion of the tissue had been removed with a straight cut, which created a structural discrepancy between the histological image and the tissue block photograph. We tried changing the resolution steps, as well as the amount of regularisation of the non-linear registration step, but ultimately the problem could only be reliably resolved by masking out the extra tissue from the target image using a hand-drawn mask. Using the mask, the registration produced excellent results with the default set of parameters and 0.8 regularisation weight.

We also noticed that the block with the anterior limb of the internal capsule had less salient anatomical features than other samples. While the orientation of other samples was correctly identified by a quick 4-direction rotation search, the registration of this sample did not succeed until a full search with 10° increments was conducted, therefore we strongly suggest adhering to this stricter routine for improved robustness.

### 3.2. Stage 2 results: Tissue block to brain slice photograph

One particular observation that we made at this stage is that most blocks with sufficiently salient anatomical features (5 out of the 6 tested) could be equally well registered using the computationally less expensive SSD cost function and the unconstrained NEWUOA optimiser instead of the standard *SSD*_*MIND*_ and BOBYQA that we described in *section 2.7*. Therefore, in Figure 10A we show the counterexample where the relative absence of anatomical features led to overscaling with these settings, and the improved results using constraints are shown in Figure 10B. As long as parameters are unconstrained, and the background area of the block photograph is masked, downscaling the unmasked area is a trivial solution for the optimiser to reduce the cost, giving rise to the risk of overscaling. In the rest of the cases this was not observed, and we attribute this to the presence of anatomical features that when mismatched between source and target, have a strong impact on the cost function, effectively constraining parameters to their optimal range. This argument is supported by the fact that using *SSD*_*MIND*_ as the image similarity metric and setting the optimisation bounds on the scale parameters to the range 0.9 – 1.1 (as described in section 2.7) led to uniform high-quality registration to the corresponding brain slice photograph in the case of all 6 blocks. (Figure 10B-E).

**Figure 10.**
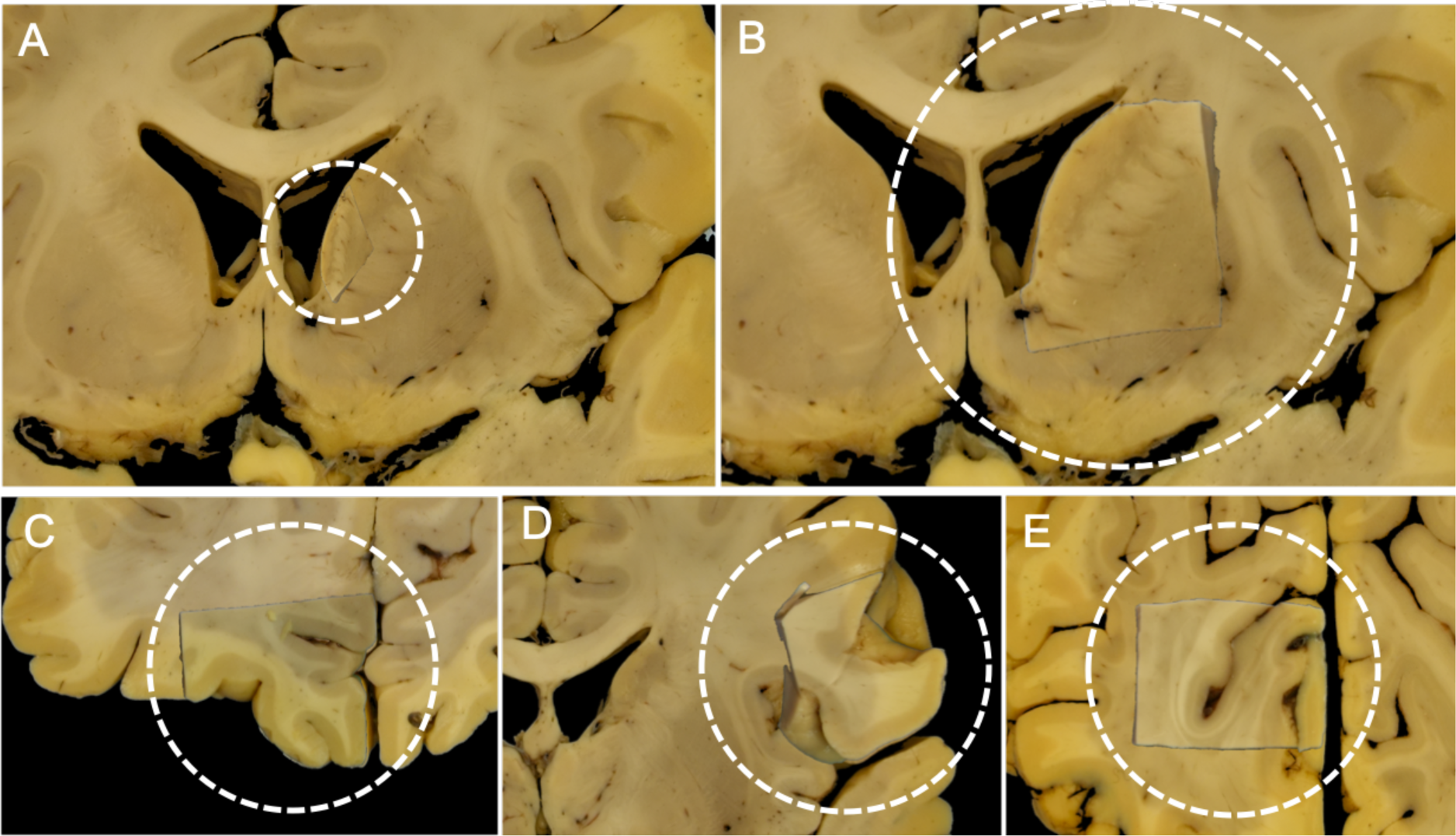
Results of stage 2 (block-to-slice registration). (A) Example of overscaling: as long as the background is masked, reducing the unmasked area is a trivial solution to reducing the cost on a tissue block that does not have enough salient features to constrain the registration (anterior limb of the internal capsule). The misregistration occurred with SSD cost and unbounded optimisation of the linear parameters. (B) Correct registration of the same block after setting the optimisation bounds on the scale parameters to (0.9– 1.1) and using MIND as the cost function. (C-E) Correct registration of various other blocks: orbitofrontal cortex, Broca’s homologue area (right hemisphere), visual cortex at the banks of the calcarine fissure.

Using these parameters, the accuracy of stage 2 was evaluated by visual comparison of the registered blocks and the underlying brain slice photographs using animations that showed both images in quick iteration. (The animations can be viewed in the Supplementary material.) We observed virtually no shift or distortion in the anatomical pattern of the blocks in the animations, indicating a degree of registration accuracy that likely surpasses the accuracy of placing landmarks to quantify the registration error. Based on our observations, the registration error incurred in this stage is negligible relative to that of the other two stages.

Possible modes of failure at stage 2 are that either (1) insertion sites are incorrectly identified on the basis of registering cut-out images, or (2) a specific tissue block is assigned to a different insertion site. Both of these would lead to catastrophic misregistration. In our experiments we never encountered a problem with the identification of the insertion sites. Even if this would happen in the future, as a last resort the pipeline allows insertion sites to be defined manually by voxel coordinates. We occasionally encountered the second problem when we used NMI as the cost, and more often when the background of the tissue block image was not masked out, or when the insertion site testing (affine registration) was performed at a coarser resolution, and the associated final cost was calculated and compared with that of other sites at full resolution. However, strict adherence to the protocol described in *section 2.7* produced high-quality stage-2 registrations for all tested brain slices (n=6). In a separate experiment we also confirmed that the stage-2 algorithm could successfully insert even as many as 6 different blocks into the same coronal slice without misregistration (images not shown).

### 3.3. Stage 3 results: Brain slice photograph to MRI volume

The accuracy of registering brain slice photographs to whole-brain MRI was tested using both simulated and real-life images.

#### 3.3.1. Experiment with simulated brain slice images

The accuracy of slice-to-volume registration was first tested on simulated data. Using TIRL, we defined a two-dimensional sampling domain in MRI space. The sampling frame was translated along the anterior-posterior axis of the MRI volume to create three sets of synthetic brain slice images at five equidistant points along the anterior-posterior axis: (1) *straight coronal slices* (no additional transformation), (2) *oblique coronal slices* (additional 3D rotation by Euler angles in the range −10°–10°), (3) *warped coronal slices* (additional *z*-axis deformations up to 6 mm according to the 2^nd^-order polynomial *P*(*x* – *x*_0_, *y* – *y*_0_), where (*x*_0_, *y*_0_) is the intersection of the slice with the anterior-posterior brain axis). Each synthetic brain slice image was subsequently registered back to the volumetric MRI data using stage 3 of the pipeline. A binary brain segmentation mask was created for each slice to define a region of interest, in which the mean registration error was evaluated by comparing the original and the registered locations of each point. Given that we register MRI to MRI in this task, the optimisation does not have to account for contrast differences between the source and target images as it would normally do. We nevertheless see this as a reasonable compromise to obtain ground truth data that the stage-3 approach can be tested against, and the results may be interpreted as ideal limits that the registration approaches with photographs or histology sections that mimic the MRI contrast.

Figure 11 shows the mean registration error after each optimisation step of stage 3. We found that the rigid and affine steps alone could register straight and oblique slices with a 0.6 mm mean registration error, which is equivalent to 1.2 voxels in MRI space. As expected for straight slices, orthogonal deformations did not make any improvement, but free-form deformation was able to take the mean registration error (0.06 mm) well below the voxel size. For warped slices, we observed a difference in the registration accuracy based on how accurate the rigid and affine stages were. In four out of five cases, the first two stages (rigid + affine) were able to achieve affine alignment with a mean registration error of 2-3 mm, which corresponds to the average deformation in these slices, suggesting that the best possible affine alignment was reached. The orthogonal and free-form registration steps both made gradual improvements, yielding a final mean registration error of 0.495 mm. In one out of the five polynomial cases, the best affine alignment could not be achieved by the linear registration steps, and the mean registration error at this stage was more than 6 mm, which would correspond to a very poorly executed brain cut. While this case shows the limits of what is achievable with stage 3 slice-to-volume registration, it is a very generous limit: if cut surfaces have elevations less than or equal to 3 mm on average, slice-to-volume registration with the presented method should be accurate to the size of a single voxel.

**Figure 11.**
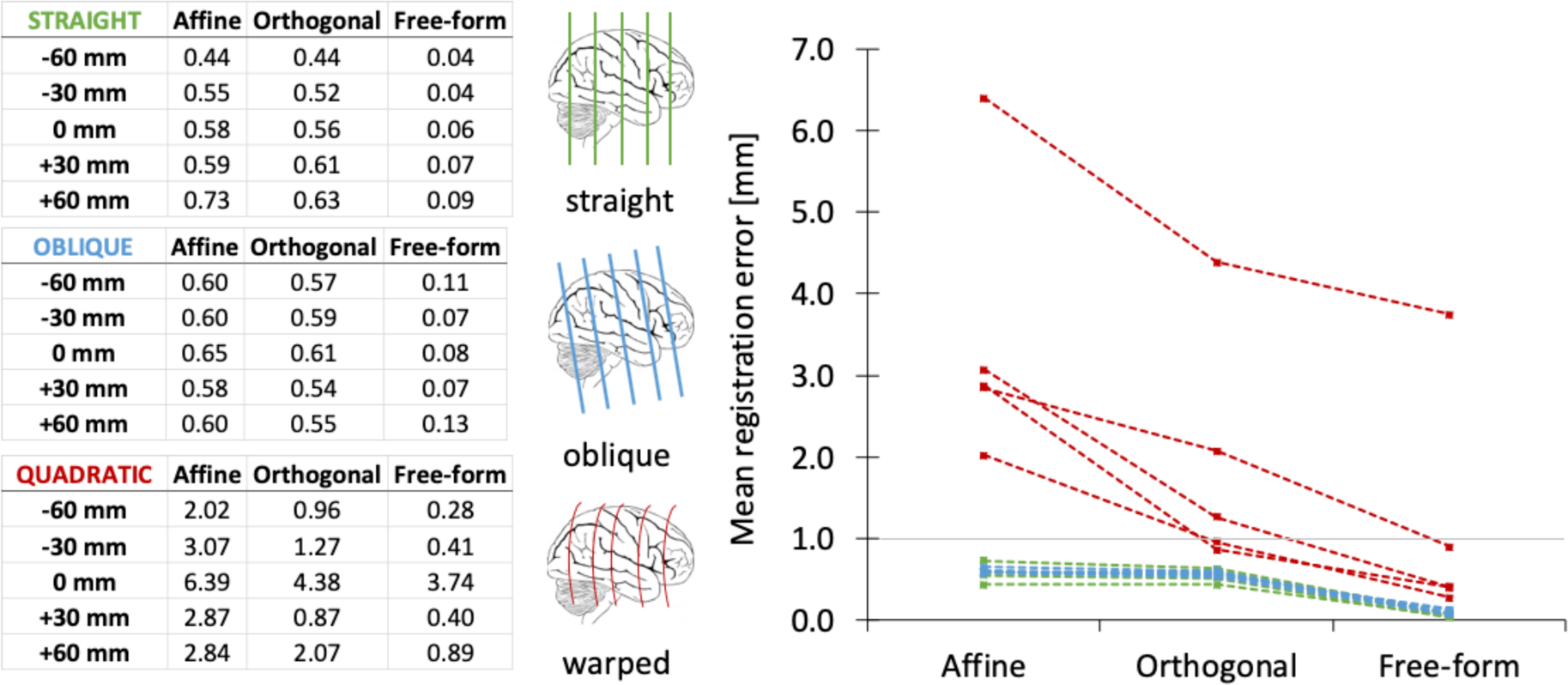
Accuracy of stage 3 (slice-to-volume registration) based on simulated brain slices. Three sets of simulated slices were created by resampling the MRI data along straight (*green*) and oblique (*blue*) coronal planes at 5 different locations along the sagittal axis, as well as slightly curved (*red*) coronal sections using 2^nd^-order polynomial transformation of the sampling domain. Consecutive optimisation steps of stage 3 gradually improved the alignment in all cases, leading to sub-millimetre final registration errors in all but one case. The one case that could not be registered by stage 3 had initial deformations larger than 6 mm, corresponding to a very poorly executed brain cut.

#### 3.3.2. Experiment with real brain slice images

To test whether our method can achieve similar registration performance on real-life images as well, we registered 5 coronal brain slice photographs, and visualised the accuracy of the registration in two complementary ways. First, we wanted to know whether in-plane deformations can be accurately compensated. Figure 12 shows a representative result with the manually segmented grey-white matter boundary of the brain slice photograph overlaid on the registered and resampled MR images after each step of the stage-3 slice-to-volume registration. The registration in this particular case was further complicated by damage to the coronal slice (Figure 12A, *asterisk*), as it was photographed after parts of the primary motor and sensory cortices had already been removed. We found that the contours were generally well-matched by the linear steps, with the largest offsets seen in the regions corresponding to the left and the right lateral sulci and temporal lobes. The orthogonal deformation step introduced a curvature of the brain slice along the left-right axis (Figure 12F), effectively shifting the cross section of an adjacent gyrus out from what is seen as subcortical white matter of the right hemisphere (right-hand side) in the photo (Figure 12B-C, *yellow arrow in the top row*), as well as fixing the alignment of the left hippocampus (Figure 12B-C, *yellow arrow in the inset*). While small, these changes are the most important from the perspective of a quantitative analysis: a registration method that had not corrected for out-of-plane deformations would have led to accidentally comparing quantitative data between grey and white matter in these regions. As an unwanted consequence of introducing slice curvature, the right hippocampal region was slightly shifted off the cutting plane by 1 voxel. (Later in *section 3.5* we will introduce a post-hoc adjustment stage to compensate for local offsets like this.) After the free-form deformation step we observed almost perfect alignment of the contours, with the largest misalignment being 0.3 mm (Figure 12D, *purple arrow*). The quantitative deformation maps in Figure 12E-G show that after all registration steps, the magnitude of in-plane deformations was on the order of 2 mm, whereas out-of-plane deformations were on the order of 2-4 mm for the majority of the slice area. The largest out-of-plane deformations (4-6 mm) were seen around the damage to the slice, but our method was able to effectively compensate for these as well. According to the Jacobian map, transformations were diffeomorphic with a maximum of 8% dilation/shrinkage of the pixels.

**Figure 12.**
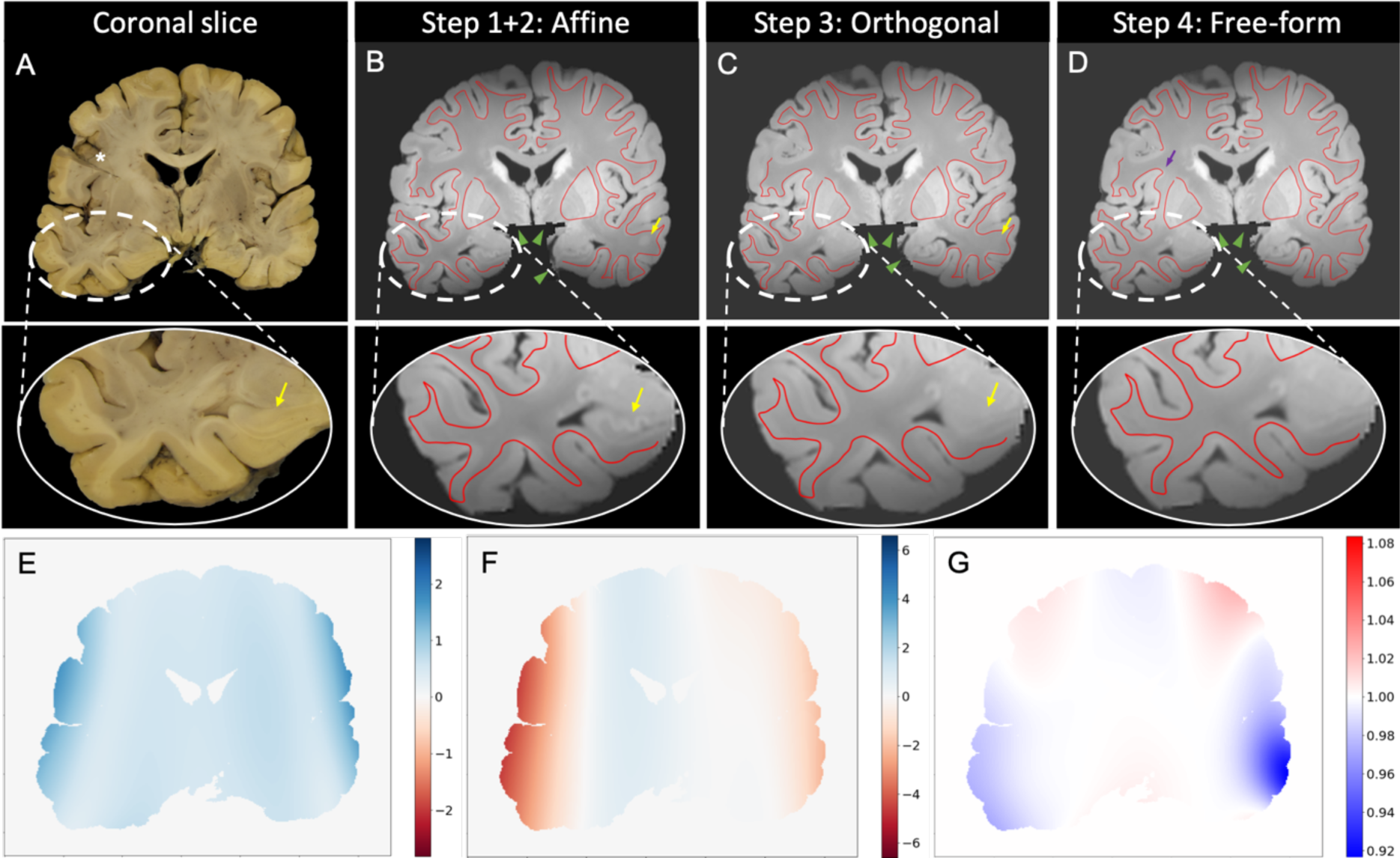
Result of stage 3 (slice-to-volume registration) on an actual brain slice image (not simulations). **(A)** Reference image used as a target of slice-to-volume registration (posterior view). The *asterisk* highlights an area where the slice was damaged due to advance resection of the primary motor and sensory cortices. The *yellow arrow* points at the cross section of the left hippocampus that will be misaligned after affine registration. **(B)** Correspondence between grey-white matter contours of the brain slice photograph (*red curve*) and the resampled MR image after affine alignment. While most of the contours match, as expected for a planar cut, large misalignments are seen in the region of the temporal lobes (*white dashed ellipse*). The *green arrowheads* point at the artificial boundaries as a result of removing the cerebellum by BrainSuite 18. The *yellow arrows* highlight regions that are misaligned after affine registration due to off-plane distortions of the slice relative to the cut surface: the left hippocampus, and a cross section of a gyrus in the subcortical white matter in the right temporal lobe. **(C)** Successful correction of the gyral and left hippocampal cross sections after the orthogonal deformation step (*yellow arrows*). In-plane deformations of the temporal regions are not yet compensated (*inset*). **(D)** Successful compensation of in-plane deformation after the final free-from registration step. The *purple arrow* shows the largest misalignment, measured as 0.3 mm. **(E)** Final in-plane deformations of the slice, showing a typical range of 0-2 mm. **(F)** Final out-of-plane distortions, which are seen to be as high as 4-6 mm, especially where the brain slice is damaged. The curvature of the slice is very prominent along the transverse (left-right) direction. **(G)** Jacobian map showing diffeomorphic transformations after final step with ±8% shrinkage/dilation of pixels.

Beyond comparing the alignment of grey-white matter contours in the registration plane, we wanted to characterise how accurately our method was compensating out-of-plane distortions of the brain slice, as this has not been addressed by previous literature. For one of the slices, we manually annotated 20 anatomical features in MRI space that visually corresponded to the anatomical features in the respective brain slice photograph and measured the distance of these points from the registered slice in MRI space. The measured distances followed a chi-distribution with a median of 0.93 mm and an interquartile range of 0.37 – 1.31 mm. This result should be interpreted with care, as the reliability of the annotation cannot be guaranteed in certain regions of the brain, where the anatomy is fairly consistent across several consecutive slices. Precise annotation in these regions requires experience and also carefully choosing the slicing orientation of the MRI volume. In our experience, even a rotation as small as 10° about one of the axes was enough to render the observable anatomy visibly very different from what was depicted in the slice photo, making the annotation process consequently very difficult. In this particular experiment, most error readings had sub-millimetre magnitude, except for 8 of the 20 that were larger than 1 mm.

To better understand the source of the registration error around these manual landmarks, we carefully inspected the registration result in these regions. The coordinates of the manual landmarks were fed into the stage-3 interpolator (as if they were the control points) to reconstruct a curvilinear slice from the MR volume (“manually registered slice”). Corresponding regions of the manually and the automatically registered MR slices were visually compared with the original slice photograph where the apparent registration error was large (Figure 13). Surprisingly, at nearly all of these locations (6 out of 8) the automated registration method was more accurate than manual annotation. This finding is important, because it shows that manual MRI slice matching by visual comparison with a 2D image is not accurate, yet it is seen as common practice where suitable software/hardware solutions for accurate MRI-histology registration are not readily available. Counterexamples (shown in the supplementary material), where the accuracy of the automated method was inferior to that of the manual landmarks were exclusively found in two cases on the edge of the brain. One of them was in the proximity of the damaged area, and the other was in a region where the pial surface was visible beneath the cutting plane (“side surface” of the brain slice) and locally biased the registration towards larger out-of-plane distortions. The accuracy of slice-to-volume registration in these regions could therefore benefit further from segmenting and masking side surfaces in brain slice photographs.

**Figure 13.**
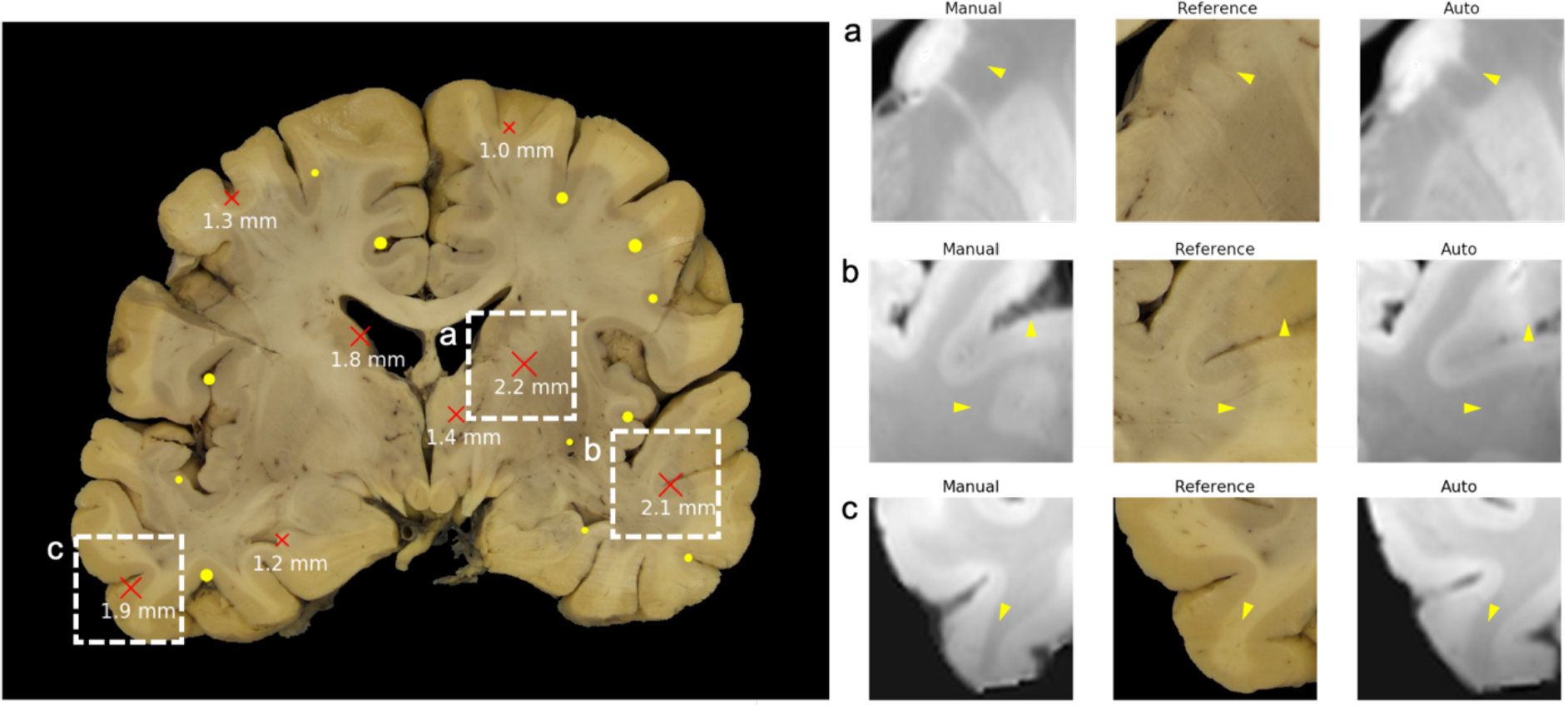
Comparison of stage-3 registration result with registration by manual landmarks. ***Left*:** Manually annotated MRI landmarks projected to the brain slice photograph. The size of the markers is proportional to the distance of the landmarks from the cut surface estimated from slice-to-volume registration by stage 3 of the pipeline. Landmarks shown as *yellow dots* are within 1 mm proximity of the surface, landmarks shown as *red crosses* are further away. Distance values for the latter are shown in millimetres. ***Right*:** Visual comparison between 2D MRI reconstructions around the manual landmarks, the reference image, and the result of stage-3 registration at three different positions (*a, b, c*) within a single slice. Careful inspection of the reconstructed MRI images reveal that the automated result is more accurate (*yellow arrowheads*), therefore the measured large distances are more indicative of annotation error than registration error, due to ambiguities in slice depth localisation.

#### 3.3.3. Slice-to-volume registration of damaged brain slices

After testing stage 3 on 5 slices, we successfully ran it on a total of 143 slices from 15 brains with identical high-quality results. The few occasions when the automatic slice-to-volume registration failed was due to some form of extreme structural discrepancy between the slice photograph and the MRI, which include: (1) significant amounts of missing tissue (cerebellum, M1S1 tissue block) or extra tissue (e.g. dislocated choroid plexus), (2) visible cortical or ventricular surfaces in the slice photograph beneath the cutting plane (“side surfaces” of the coronal brain slice), and (3) large local displacements such as the closing or the opening of the interhemispheric fissure as a result of one hemisphere moving toward or away from the other one.

In all cases, the problem of missing tissue was successfully addressed by creating hand-drawn masks (Figure 14B) for the target image (slice photo), which recovered the registration accuracy for most of the unmasked regions, but lead to larger deviations closer to the masked region due to lack of supporting features (Figure 14). We found that the problem of side surfaces could be most effectively addressed by taking photographs of both sides of the brain slices and registering the one with less side surfaces visible. Alternatively, masks can be generated automatically for side surfaces by affine registering the images of adjacent slices that display the same cut surface, segmenting non-matching regions and adding them to the target mask. The problem of hemisphere separation only affected a few slices in our case, and we resorted to registering hemispheres separately in these cases.

**Figure 14.**
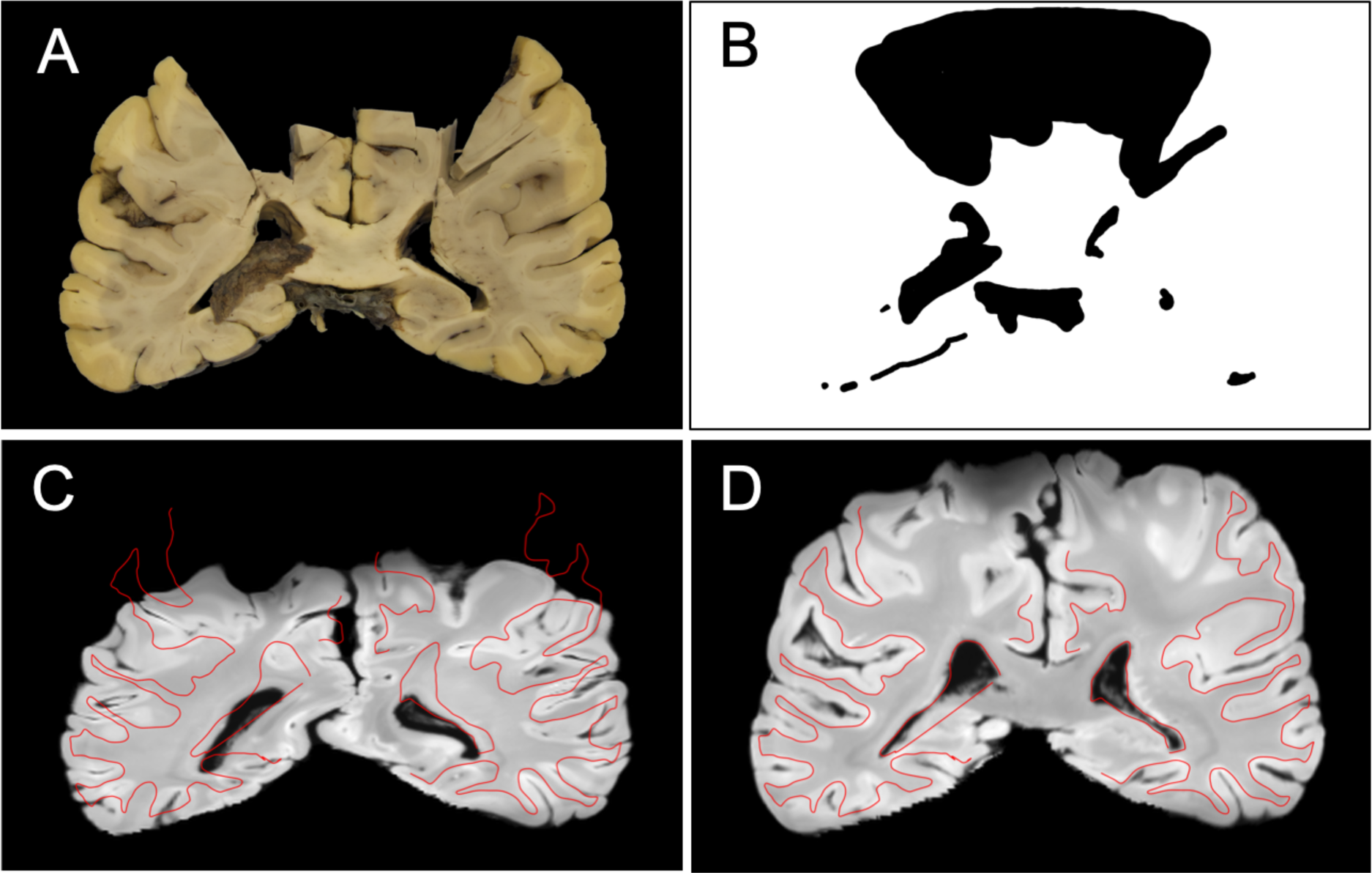
Result of slice-to-volume registration of a severely damaged coronal brain slice. **(A)** Coronal brain slice photograph with bilateral hiatus in the sensorimotor regions. **(B)** A hand-drawn binary mask for cost-function weighting. **(C)** Registration result without using the target mask. The *red curve* is an overlay of the manually segmented grey-white matter contour of the brain slice photograph. **(D)** Registration result with the hand-drawn target mask. The accuracy of the corrected registration is qualitatively similar to that on non-damaged slices, but misalignments are slightly larger in the proximity of the masked regions due to the relative absence of driving features.

### 3.4. Combining stages 1-3: histology-to-MRI registration

To achieve end-to-end histology-to-MRI registration, we combined the histology-to-block, block-to-slice and slice-to-volume registration stages according to the halfway method as described *section 2.9*. Figure 15 shows a representative final result of the registration between MRI and histology for five of the six blocks stained for ferritin. Qualitatively identical registrations were obtained with the PLP stains of the same five blocks. A three-dimensional rendering of the registered histological sections can be seen in Figure 16.

**Figure 15.**
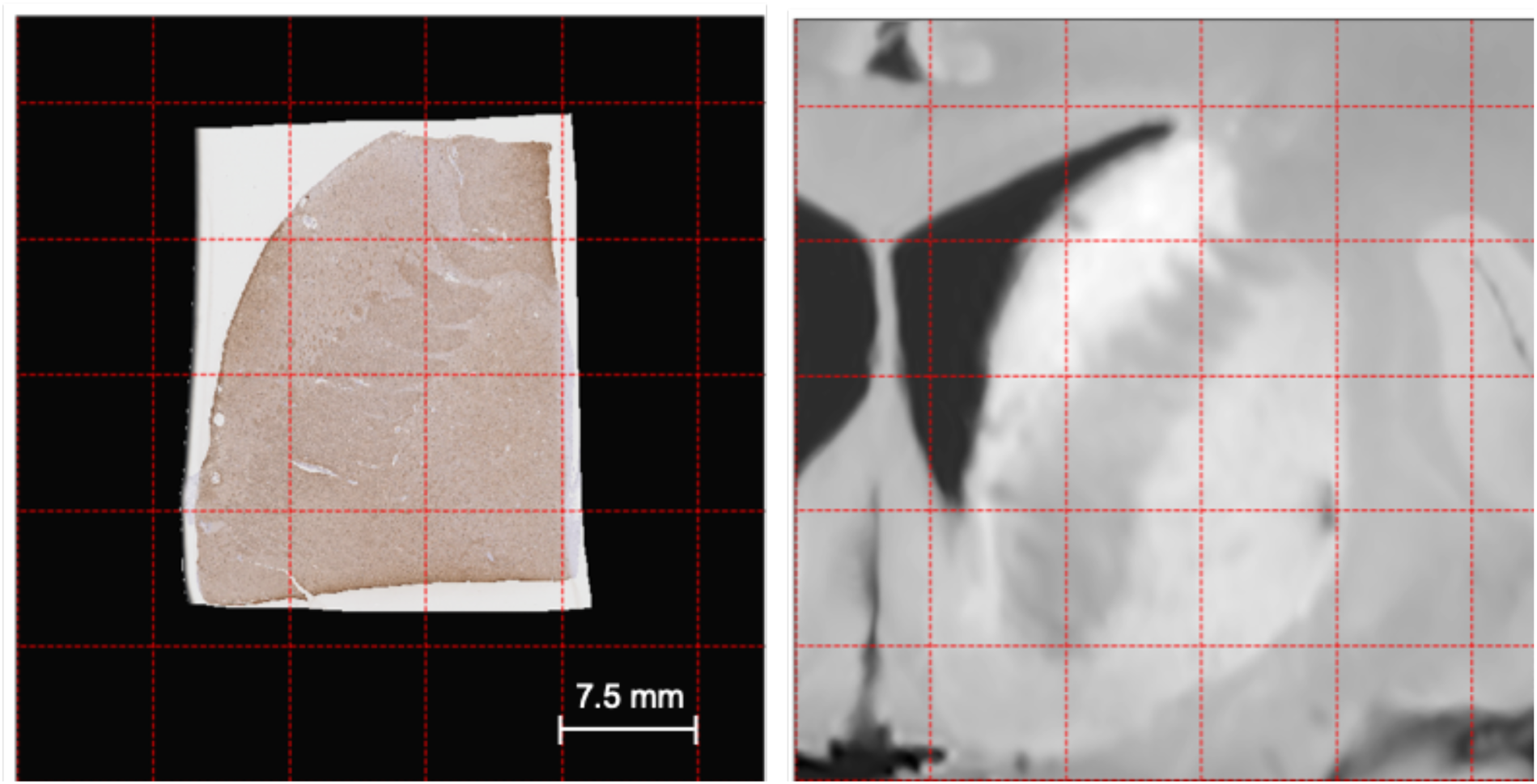
End-to-end histology-to-MRI registration by combining stages 1-3. ***Left:*** Histological section of the anterior limb of the right internal capsule stained for ferritin. The image was resampled at the resolution of the tissue block photograph (50 μm/pixel). ***Right:*** The corresponding 2D section of MRI resampled at the resolution of the tissue block photograph. The *red gridlines* are provided as a common spatial reference for comparing the images.

**Figure 16.**
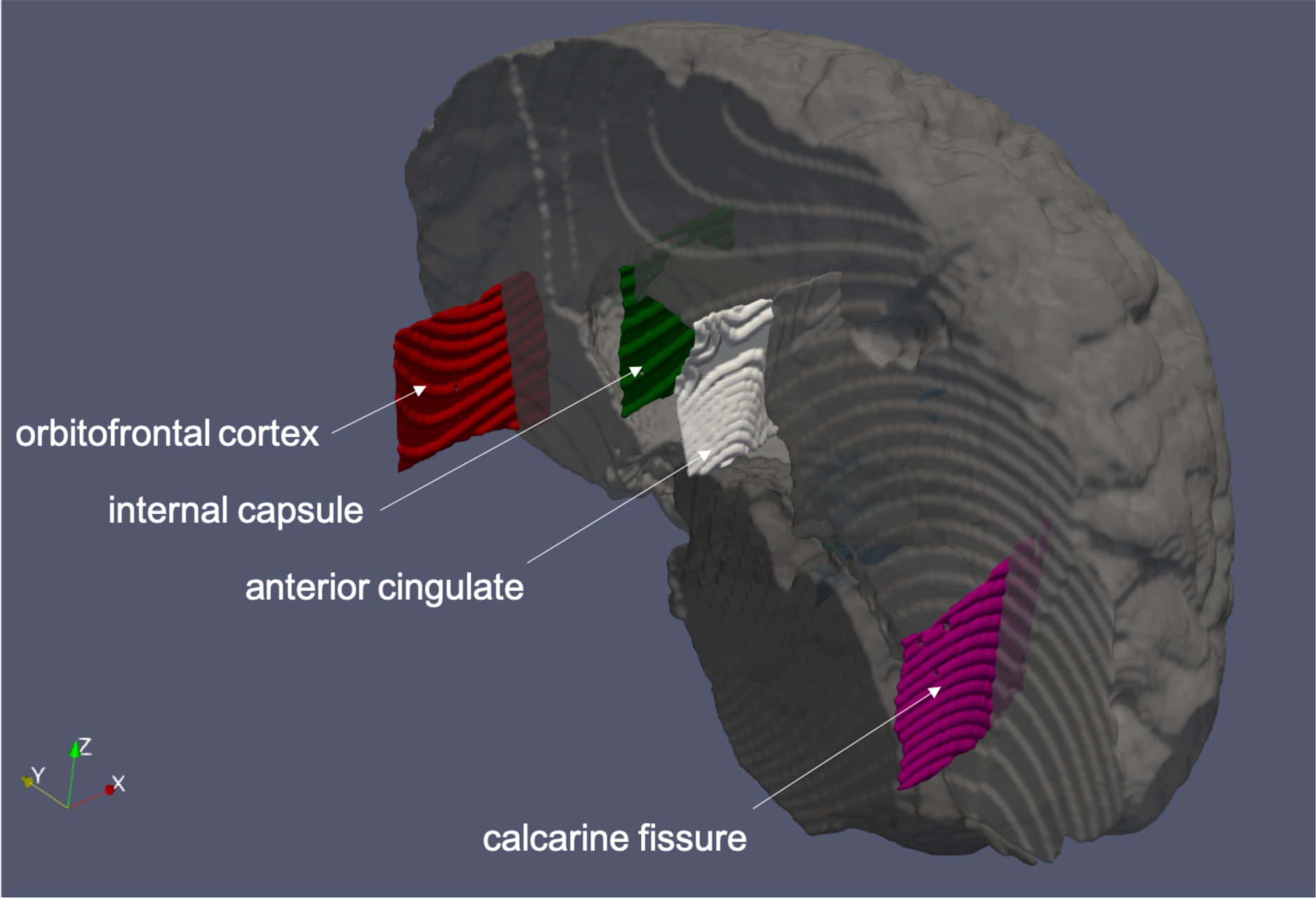
Three-dimensional model of the post-mortem brain showing a subset of the registered histological sections. The registered sections are represented by their curvilinear image domains, which are larger than the actual sections. The left-hand side of the brain was removed for better visualisation, and two registered sections are not visible on this median sagittal surface view. Note that geodesic lines are accented because the surfaces were reconstructed from voxel-wise labels in MRI space (voxel size: 0.5 mm).

In the case of a single tissue block, which was sampled symmetrically to the mid-sagittal plane to contain the cross section of the corpus callosum and the anterior portion of the cingulate gyri from both hemispheres, we noticed that both the ferritin and the PLP stains registered imperfectly with the MR volume. The error was confined to a region within the image where one of the gyri had a significantly larger separation from the corpus callosum in the MRI image, that was not compensated by the free-form deformations of stage 3 (slice-to-volume registration). Large local deformations of this kind are typically challenging because they are heavily penalised by membrane energy regularisation, and only a condensed set of local control points could accurately represent them without affecting the alignment in more distant regions of the image. While the current implementation achieves sufficient accuracy in the largest portion of this image, we anticipate that the observed type of registration error may be better addressed in future versions of TIRL by suitable changes to stage-3 registration, as explained in the *Discussion* section.

### 3.5. Stage 4 (optional): refinement by direct histology-to-MRI registration

In the two cases where an LFB+PAS stain was also performed subsequent to all other stains, we noticed that the anatomical consistency between the histological images and tissue block photo was not perfect due to the slicing depth problem (Figure 17A). As tissue blocks are embedded in paraffin, which will generally have a slightly larger volume than the block itself, it cannot be guaranteed that the surface of the paraffin block is parallel to the surface of the tissue block. For sectioning in a microtome, the blocks are trimmed to remove any excess paraffin from the surface of the block to fully expose the tissue. During this process some sections come off the block as partial sections and are therefore discarded. Depending on the angle of sectioning, the first full slice of tissue may come from as deep as 0.5–1 mm (corresponding to an angle of 2° for a 30 mm long block). In the case of multiple stains, or when stains need to be repeated for quality reasons, this problem is further exaggerated: the deeper the block is sampled, the less consistent the stained histological sections will be with the surface anatomy of the blocks as seen in the photographs. This means that the inaccuracies at stage 1 (histology-to-block registration) should be dealt with less aggressively, as they may reflect true differences between the stained section and the photograph. Instead a higher regularisation weighting is preferred in these cases, to preserve the structural self-consistency of the histological section while compensating for some of the distortions.

**Figure 17.**
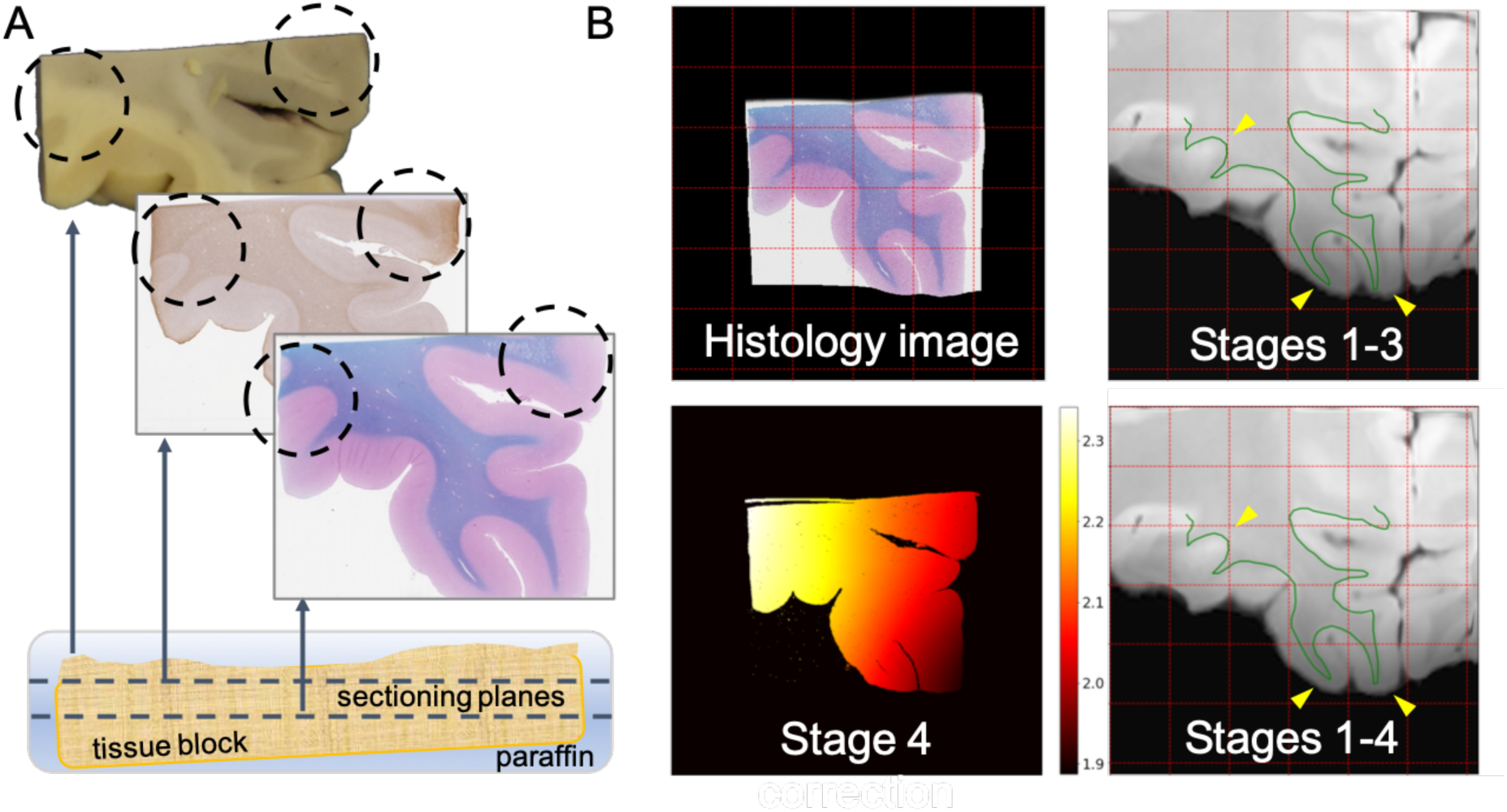
The slicing depth problem and the optional 4th stage of the pipeline (direct histology-to-MRI registration). (A) The slicing depth problem. If the block is slightly tilted relative to the surface of the paraffin embedding, a portion of the surface is abraded by the microtome before the first full section of tissue is obtained. Subsequent slices are sampled at a relative depth from the surface, which may cause substantial anatomical discrepancy between the surface of the block (as seen on the photograph) and the surface of the slide. Consequently, the final end-to-end histology-to-MRI mapping will be inaccurate. **(B) Stage-4 registration.** *Top row*: Alignment of the LFB+PAS-stained section (*top left*) of the left OFC with the MRI before (*top right*) stage-4 correction. The *green curve* is the overlay of the hand-drawn grey-white matter boundary of the transformed histological image. *Bottom row*: Result of stage-4 slicing depth correction shows as much as 0.4 mm elevation difference across the surface of the section, corresponding to a 1° tilt. (The numbers in the colour bar represent distance along the *z*-axis in millimetres after stage-4 registration starting from the best 3D affine alignment that was inferred from the combination of the previous three stages.) *Bottom right*: Improved alignment (*yellow arrowheads*) of the same section with MRI after stage-4 correction.

To compensate for the depth problem, in these two cases an additional fourth stage of the pipeline was introduced. Stage 4 aims to fine tune the alignment between the histological section and the MRI image by performing a direct registration between the MRI data and the histological section after the latter is initialised to MRI space by the three main stages. First, the histological section was resampled on the intermediate domain (tissue block photograph) using the transformation chain from stage 1. The optimised transformation chains from the second and third stages were then concatenated and attached to the domain of the resampled histological image, mapping it into MRI space. The free-form deformation object from stage 3 was redefined within the combined transformation chain such that its new control points were concentrated on the area of the histological image instead of being scattered across the whole coronal brain slice. The parameters for this new transformation were fitted to preserve the previously optimised in-plane and out-of-plane deformations within the area of the resampled histology image. The updated chain of transformations was applied to obtain the physical (MRI-space) coordinates of the resampled histological image. The centre of the inserted histological section was determined by averaging the physical coordinates, and the normal vector of the histological image was calculated as the 3^rd^ principal component of the physical coordinate array. The sample was gradually shifted in MRI space along the normal vector in the range −2.5–2.5 mm, while the 3D rotation parameters were optimised within 12° in both directions from the initial values for minimum *SSD*_*MIND*_ cost using the BOBYQA optimiser. Finally, starting from the best position and rotation of the histological image, orthogonal and later free-form deformations were optimised with membrane energy regularisation within 1 mm of their initial values to obtain the final registration between histology and MRI. Figure 17B shows the alignment of the more offending LFB+PAS stained section before and after the stage-4 correction. The correction included shifting the histological image from its original position by approximately 2 mm and introducing 0.4 mm through-plane deformation.

## 4. Discussion

In the past three decades a handful of studies have addressed different aspects of registering histology and MR images by semi-automatic methods. However, most of these algorithms were tailored to a specific application and/or they were implemented as in-house scripts, which are no longer accessible to the larger community. In comparison, 3D-to-3D image registration is a fundamental operation in the field of neuroimaging that most higher-level analysis methods depend on. Consequently, 3D image registration tools are well-established and lie at the core of popular analysis toolboxes, such as FSL, SPM, FreeSurfer, BrainSuite, *etc*. On the contrary, similar registration tools are less well developed and mostly non-existent for hybrid MRI/histology datasets, which has precluded the evolution of equally powerful analysis toolboxes for this kind of data. Due to time and labour constraints, neuropathology facilities are collecting the overwhelming majority of their histology data in the format of stand-alone histological sections, not 3D stacks. The alignment of these images with volumetric MRI data is a tedious and imperfect manual process, which obviates bias-free quantitative analysis, and limits the number of samples and subjects that can be studied at once. With limited sample sizes and imperfect matching, studies that aim to analyse MRI signal changes in diseased tissue may not capture the significant interindividual variations in the spatial and temporal extent of a disease (which are recognised as different phenotypes in neurodegenerative conditions). Consequently, slice-to-volume histology-to-MRI registration is a fundamental operation that must be automated before higher-level analyses can be performed on large volumes of this type of data, and stable conclusions can be made about the pathological interpretation of characteristic MRI signal changes.

In this paper, we presented an automated registration pipeline for sparsely sampled histology data and post-mortem MRI. Our method does not require specialised cutting or stain automation hardware for tissue processing and reduces the imperfections of alignment that arise from freehand brain cutting, which altogether make it suitable for integration into routine neuropathological practice. The first three stages of the pipeline support full automation of the registration, provided that suitable dissection photographs are available. Otherwise, the optional stage 4 may be used on its own as a semi-automatic tool to register histological sections to volumetric MRI after manual initialisation, although this feature should be tested more thoroughly. Most importantly, all stages of the pipeline are embedded in the more general open-source (Python 3.7) framework, TIRL, that allows them to be modified for a wider range of applications, potentially including small-animal and non-human primate neuroimaging, as well imaging other organs and tumours. Finally, we have decided to include TIRL and the pipeline in FSL to facilitate continuous improvement to the framework and the registration techniques therein, as well as to encourage the development of further analysis tools for hybrid MRI/histology datasets.

As with all methods, our pipeline also has certain limitations. First, while we committed significant efforts to ensure that the pipeline can perform all stages automatically, this is subject to a set of assumptions about the input data. Based on the conditions under which the pipeline was tested, we recommend observing the following precautions:

1. Histological sections should be sampled close (<2.5 mm) to the surface of the tissue blocks. Care should be taken to avoid staining artefacts and tears during the sectioning process. Stains with grey-white matter contrast must be used for registration.
2. The approximate location and rough orientation of coronal sections must be known in advance.
3. Photographs should be taken at high resolution, under diffuse lighting conditions, on a clean, matte surface that has a distinct colour from the brain tissue. Brain slices should be photographed on both sides avoiding glares. The approximate mm/pixel resolution of the photographs should be recorded.
4. MRI should be acquired at high resolution (0.5-1 mm) with sufficient grey-white matter contrast. For post-mortem imaging, formalin-fixed brains should be immersed in an inert fluorocarbon medium (e.g. Fluorinert) to minimise the background signal, and scanned in a suitably shaped plastic container to prevent large deflections of the hemispheres, the brainstem and the cerebellum.

While the above prescriptions may seem very restrictive, they directly reflect our own experimental approach that was used to test both TIRL and the pipeline. We strongly believe that the capability of the software tools that were developed for this project extend beyond the scope of the current application, and the flexibility of TIRL allows many of the above restrictions to be loosened.

Registering histological stains with little or no grey-white matter contrast is beyond the scope of the current work, and is therefore not readily supported by the current pipeline. However, the results of a recent grand challenge competition (ANHIR) [66] might be used in the future to register histological sections with different stains in advance, and the ones with appropriate grey-white matter contrast to the MRI. Alternatively, these images could be registered linearly by matching outer contours or non-linearly by manually defined landmarks. Either of these approaches would be a straightforward extension to the current cost and transformation libraries of TIRL.

Generality and optimal computational performance are often competing demands in software engineering. Several features have been implemented in TIRL to make computations more effective, such as parallel processing, chunked interpolation, function caching, optimising sub-chains of linear transformations by affine replacement, and avoiding interpolation of displacement fields where the field is defined over the same domain as the image. That said, greater emphasis was put on preserving the generality of the framework. Therefore, some of the computations may benefit from further optimisation, which lie beyond the scope of the current work. One particular improvement would consider adaptive control point placement in stages 3 and 4. Instead of initialising a fixed set of control points and optimising the corresponding deformation parameters all-at-once, one could start with a smaller set of control points and gradually increase their count. Whenever sufficient convergence is reached with the current set, a new control point would be added where image dissimilarity is the greatest. This strategy would provide better control over large local displacements by permitting local clusters of control points, altogether leading to fewer registration errors.

Our experiments were carried out on a MacBook Pro computer with a dual-core 2.7GHz CPU and 8 GB of RAM. The typical runtimes were ∼2 minutes for stage 1, ∼30 minutes for stage 2 (with 6 insertion sites), 1-2 hours for stage 3 (using 50 control points), and ∼15 minutes for stage 4 (where needed). For relatively undistorted slices, it is possible to reduce the runtime of stage 3 by using fewer (e.g. 16 or even less) control points instead of 50. Running the stages in parallel can also save significant amounts of time. With the adaptive control point placement described above, stage 3 and 4 could benefit from faster convergence, as only a subset of parameters would need to be optimised in the first iterations, and the runtime of these stages could be consequently greatly reduced.

Despite the current limitations, our method allows automated registration of histology to MRI without labour-intensive sequential sampling and volumetric reconstruction of histology as opposed to the majority of existing methods. Contrary to the methods of *Kim* et al and *Singh* et al for slice-to-volume registration, our method does not require manual intervention for the majority of the cases and uses more precise local deformations by radial basis functions instead of polynomial transformations. Extending the framework-building approach of *Osechinskiy* et al, using TIRL we successfully applied the MIND cost function [43] to register not only hemispheres, but whole brain slice photographs as well as small histological samples that could otherwise not be directly registered to MRI. Most importantly, our results demonstrate that histological sections are not immune to out-of-plane deformations due to free-hand cuts through the brain. Nevertheless, using TIRL, it is possible to align these images with MRI data with sub-millimetre precision, which has important implications for biomarker research.

Establishing novel imaging-derived biomarkers that can sensitively and specifically indicate the presence of a disease is one of the chief goals in modern medical imaging. Classic radiological signs such as signal hypo- and hyperintensities in weighted MRI scans have suboptimal disease specificity due to the complex dependency of the MRI signal on both the acquisition parameters and a spectrum of elementary disease-related changes in tissue microstructure. By modelling the signal behaviour in the healthy and the diseased state of tissue, advanced microstructural MRI methods can be more specific to these elementary changes, and thus the underlying pathological process. The clinical translation of these methods requires thorough validation against histopathology, which will hopefully be facilitated by the availability of MRI-histology registration tools. A more exciting implication is that as soon as suitably large MRI/histology datasets become available, these could be used by learning algorithms to detect subtle changes of the MRI signal related to tissue pathology, which would otherwise be unnoticeable during routine radiological assessment. A new generation of such histology-inspired imaging biomarkers could be more sensitive predictors of disease. In neurodegenerative conditions, increased sensitivity to the early sub-clinical stages of the disease is critical, as the anticipated benefit from any therapeutic approach is proportional to the remaining functional capacity of the central nervous system.

## 5. Conclusion

The capabilities of a novel image registration framework, TIRL, were presented in the context of creating an image registration pipeline for post-mortem MRI and sparsely sampled histology data. Small stand-alone histological sections were successfully registered to post-mortem whole-brain MRI without manual intervention in most cases, achieving a final accuracy of 0.5 – 1 mm. In-plane and out-of-plane deformations of the sampling surface were also taken into account in the process. The method does not require additional specialist hardware for tissue pre-processing, therefore it can be integrated into routine neuropathological practice. Both TIRL and the registration pipeline is released as part of FSL, facilitating MRI-histology validation studies to be carried out in much larger cohorts than previously possible. The customisability of the presented software tools allows them to be reused in other research contexts, and hopefully provide the necessary grounds for future explorative research into a new generation of histology-inspired microstructural imaging biomarkers, that can be more sensitive predictors of neurodegeneration.

## List of abbreviations

ALS: amyotrophic lateral sclerosis,
ANHIR: Automatic Non-rigid Histological Image Registration,
BOBYQA: Bound Optimisation by Quadratic Approximation,
bSSFP: balanced steady-state free precession sequence,
CR: correlation ratio,
CT: computed tomography,
DOF: degrees of freedom,
FSL: FMRIB Software Library,
FWHM: full width at half maximum,
H&E: haematoxylin and eosin (histological stain),
LFB+PAS: Luxol fast blue combined with the periodic acid-Schiff procedure (histological stain),
MIND: Modality-Independent Neighbourhood Descriptor,
MND: motor neuron disease,
MRI: magnetic resonance imaging,
NEWUOA: New Unconstrained Optimisation Algorithm,
NMI: normalised mutual information,
OBB: Oxford Brain Bank,
OFC: orbitofrontal cortex,
PLP: proteolipid protein,
pTDP-43: phosphorylated TAR-DNA binding protein 43 kDa,
SPM: Statistical Parametric Mapping (software),
SSD: sum of squared differences,
TIRL: Tensor Image Registration Library,
TPS: thin-plate spline

## Authors’ contributions

*I. N. Huszar*: Designed, implemented, tested TIRL and all scripts of the registration pipeline, created figures, wrote manuscript.

*M. Pallebage-Gamarallage*: Designed the histopathological protocol of the MND study, dissected brains, took dissection photographs and created stained histological specimens, edited manuscript.

*S. Foxley*: Designed the post-mortem MRI protocol of the MND study and acquired MRI data, edited manuscript.

*B. C. Tendler*: Created post-processing pipeline for post-mortem MRI data, edited manuscript.

*A. Leonte*: Prepared stained histological specimens of the anterior cingulate cortex, edited manuscript.

*M. Hiemstra*: Prepared stained histological specimens of the hippocampus, edited manuscript.

*J. Mollink*: Prepared stained histological specimens of the hippocampus, edited manuscript.

*A. Smart*: Prepared various stained histological specimens, edited manuscript.

*S. Bangerter-Christensen*: Prepared various stained histological specimens, edited manuscript.

*H. Brooks*: Prepared LFB-stained histological specimens, edited manuscript.

*O. Ansorge*: Designed MND study, provided neuropathological expertise, and material from the Oxford Brain Bank, edited manuscript.

*M. R. Turner*: Designed MND study, provided neurological expertise, edited manuscript.

*K. L. Miller*: Designed MND study, provided MRI physics expertise, edited manuscript.

*M. Jenkinson*: Provided image analysis expertise, designed TIRL, the registration pipeline and the experiments, edited manuscript.

## Acknowledgement

The authors express their gratitude to the donors and benefactors of the Oxford Brain Bank, that kindly provided all human tissues for this study. Core funding for the Oxford Brain Bank was provided by the Medical Research Council (MRC), the NIHR Oxford Biomedical Research Centre and the Brains for Dementia Research programme, jointly funded by Alzheimer’s Research UK and Alzheimer’s Society and Brains for Dementia Research. INH was supported by the Engineering and Physical Sciences Research Council (EPSRC) and the MRC (EP/L016052/1), and the Clarendon Fund in partnership with the Chadwyck-Healey Charitable Trust at Kellogg College (Oxford). MJ and OA were supported by the National Institute for Health Research (NIHR) Oxford Biomedical Research Centre (BRC). MPG, SF and the dataset used in this study were funded by an MRC Project Grant (MR/K02213X/1). KLM, BCT and JM were funded by a Wellcome Trust Senior Research Fellowship (202788/Z/16/Z). The Wellcome Trust provided core funding for the Wellcome Centre for Integrative Neuroimaging (203139/Z/16/Z).

