## Supplementary material for "Tensor Image Registration Library: Automated Non-Linear Registration of Sparsely Sampled Histological Specimens to Post-Mortem MRI of the Whole Human Brain"

### Slide 1
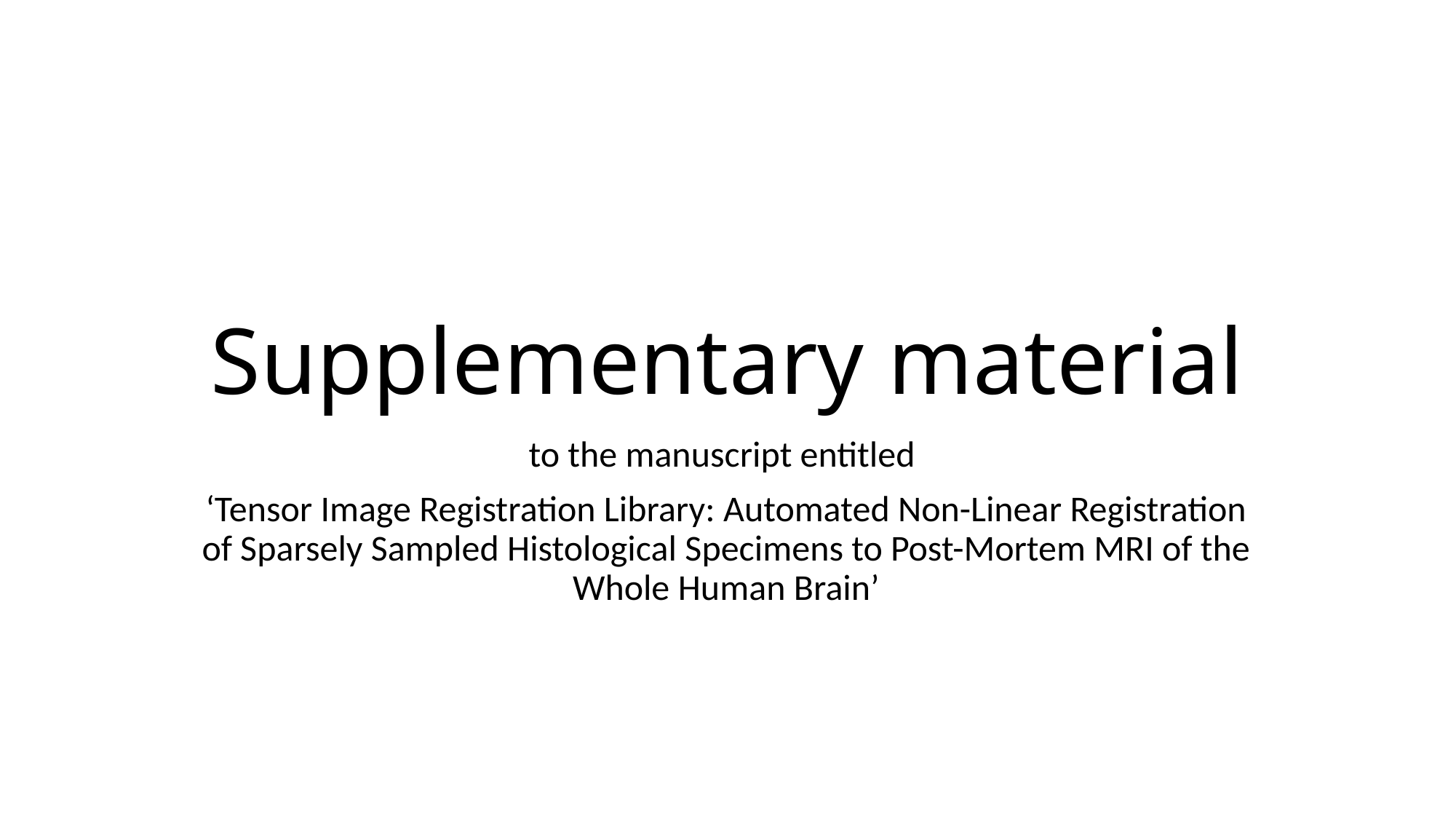

#### Supplementary material
to the manuscript entitled
‘Tensor Image Registration Library: Automated Non-Linear Registration of Sparsely Sampled Histological Specimens to Post-Mortem MRI of the Whole Human Brain’

### Slide 2
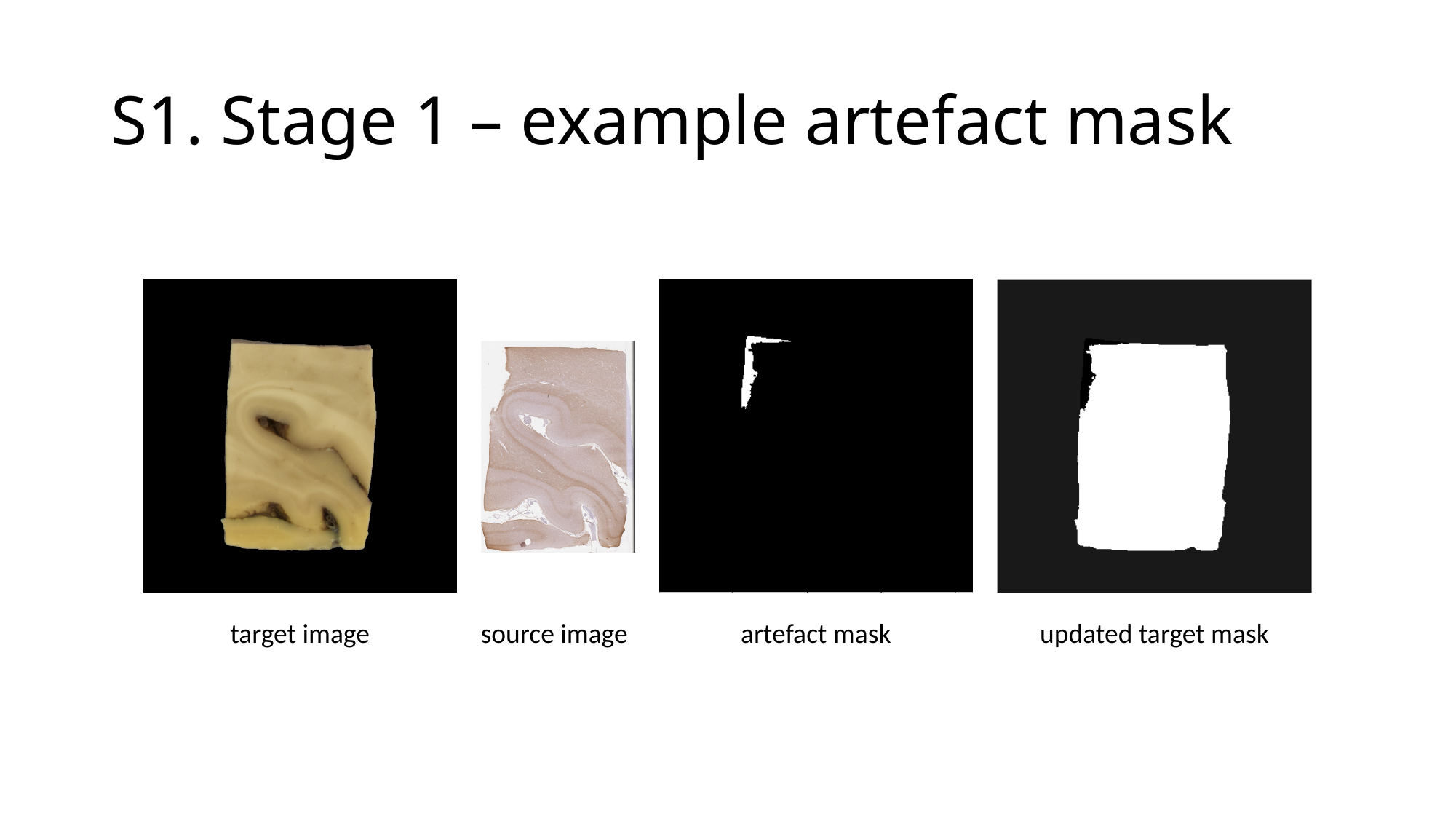

#### S1. Stage 1 – example artefact mask
target image
source image
artefact mask
updated target mask

### Slide 3
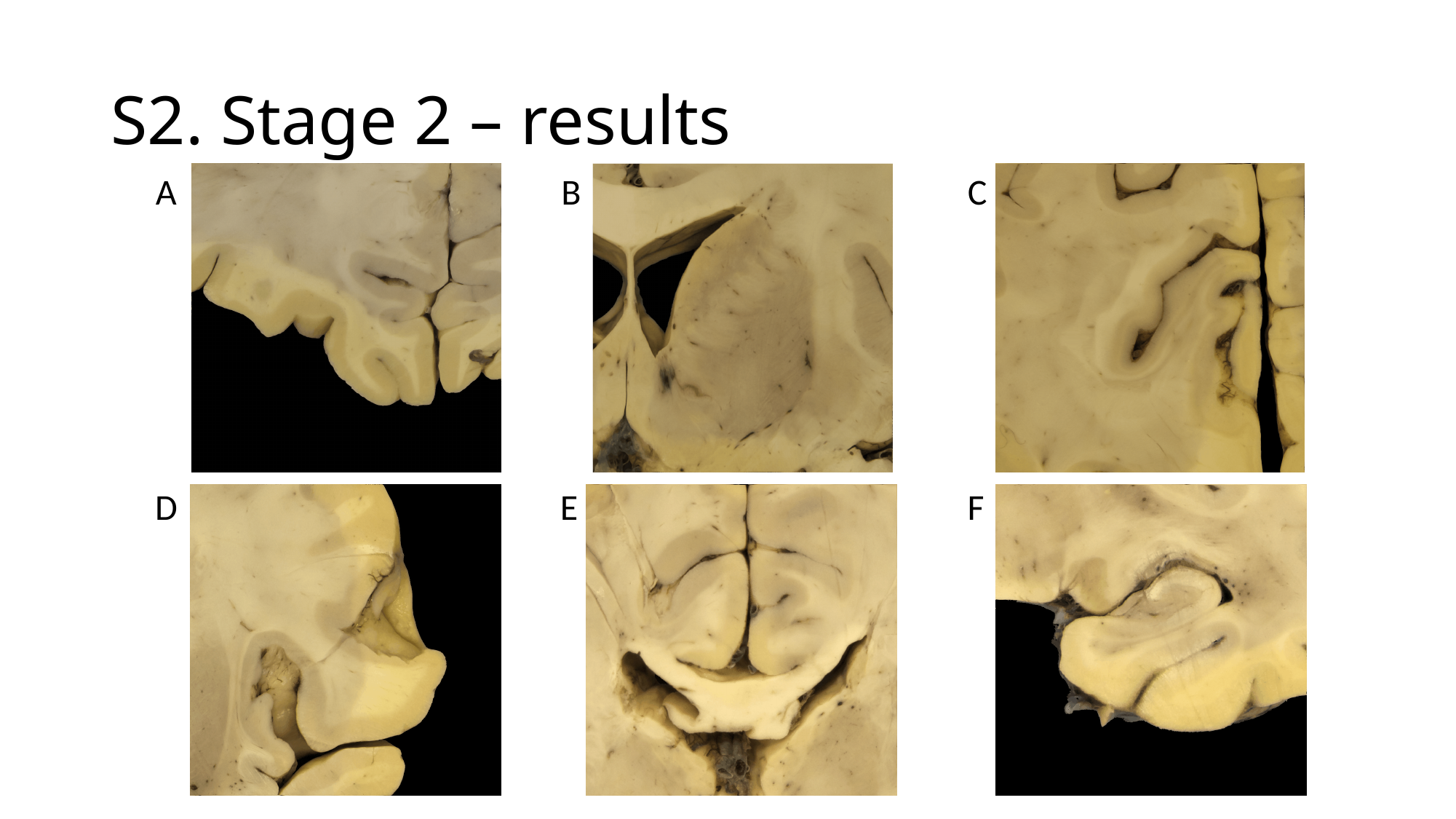

#### S2. Stage 2 – results
C
B
A
F
E
D

### Slide 4
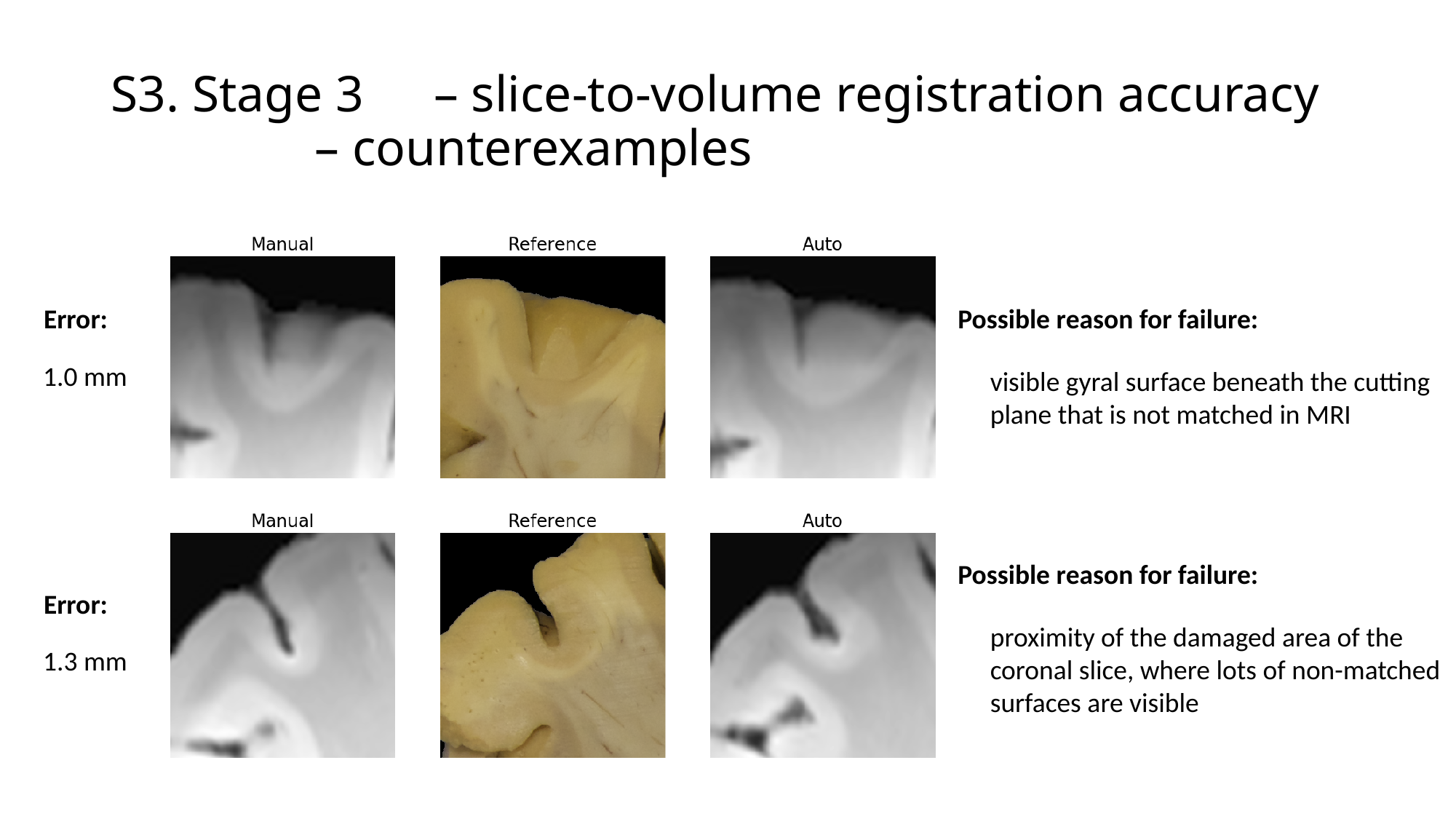

#### S3. Stage 3	– slice-to-volume registration accuracy – counterexamples
Error:
Possible reason for failure:
1.0 mm
visible gyral surface beneath the cutting
plane that is not matched in MRI
Possible reason for failure:
Error:
proximity of the damaged area of the coronal slice, where lots of non-matched surfaces are visible
1.3 mm
